# Impacts of changing ocean circulation, temperature, and food supply on larval recruitment of purple sea urchins in Southern California: A biophysical modeling study

**DOI:** 10.1101/2025.07.30.667731

**Authors:** Rachel D. Simons, Daniel K. Okamoto

## Abstract

Changes in ocean conditions among years and across decades can alter populations in marine ecosystems. This study evaluates the impact of changing ocean conditions on larval recruitment of purple sea urchins in the Southern California Bight using 3D biophysical modeling, surface chlorophyll, and recruitment data. The influence of circulation, temperature, and food supply on larval recruitment is quantified using five modeled variables, larval dispersal distance, larval source location, larval food supply, and temperature exposure for larvae and adults, which are derived for 18 years and five larval recruitment sites. Sensitivity testing of the variables to different plankton larval durations (PLDs), larval behaviors, and nearshore retention is performed. All variables are found to be relatively insensitive to changes in PLD greater than 26 days. Larval dispersal distance and source location, representing changes in circulation, are found to be more sensitive to larval behavior and nearshore retention than larval food supply and temperature exposure for larvae and adults. All variables are statistically compared to recruitment field data. Temperature exposure for adults during the fall reproductive season is found to be a strong driver of larval recruitment while temperature exposure for larvae during the spring recruitment season is not. Food supply is not found to be a driver of larval recruitment. Circulation is found to be a driver of larval recruitment if larvae have behavior that reduces their dispersal distance, allowing them to come from source sites near to the recruitment sites. Overall, we hypothesize that larval behavior which reduces dispersal improves recruitment and that the timing of recruitment and reproduction can predict the impact of temperature on recruitment.

## 1.0 Introduction

Extreme changes in ocean conditions, such as temperature, circulation, and primary productivity, pose significant threats to marine ecosystems and are predicted to alter the distribution and productivity of marine species, in part by affecting larval recruitment (Doney et al. 2012; Sydeman et al. 2014). Improving our understanding of larval recruitment can explain and predict shifts in the dynamics of populations of fished and ecologically important species (Botsford et al. 1998; Myers 1998; Pineda et al. 2010; Stige et al. 2013). Thus, quantifying how ocean conditions shape spatial and temporal variation in larval dispersal and recruitment is critical to our understanding how climate drivers influence marine populations. In this study, we explore the impacts of changing ocean conditions on larval recruitment by investigating the relationship between larval recruitment of purple sea urchins and ocean conditions of circulation, temperature, and food supply in the Southern California Bight (SC Bight, Fig. 1) using 3D biophysical modeling.

**Figure 1:**
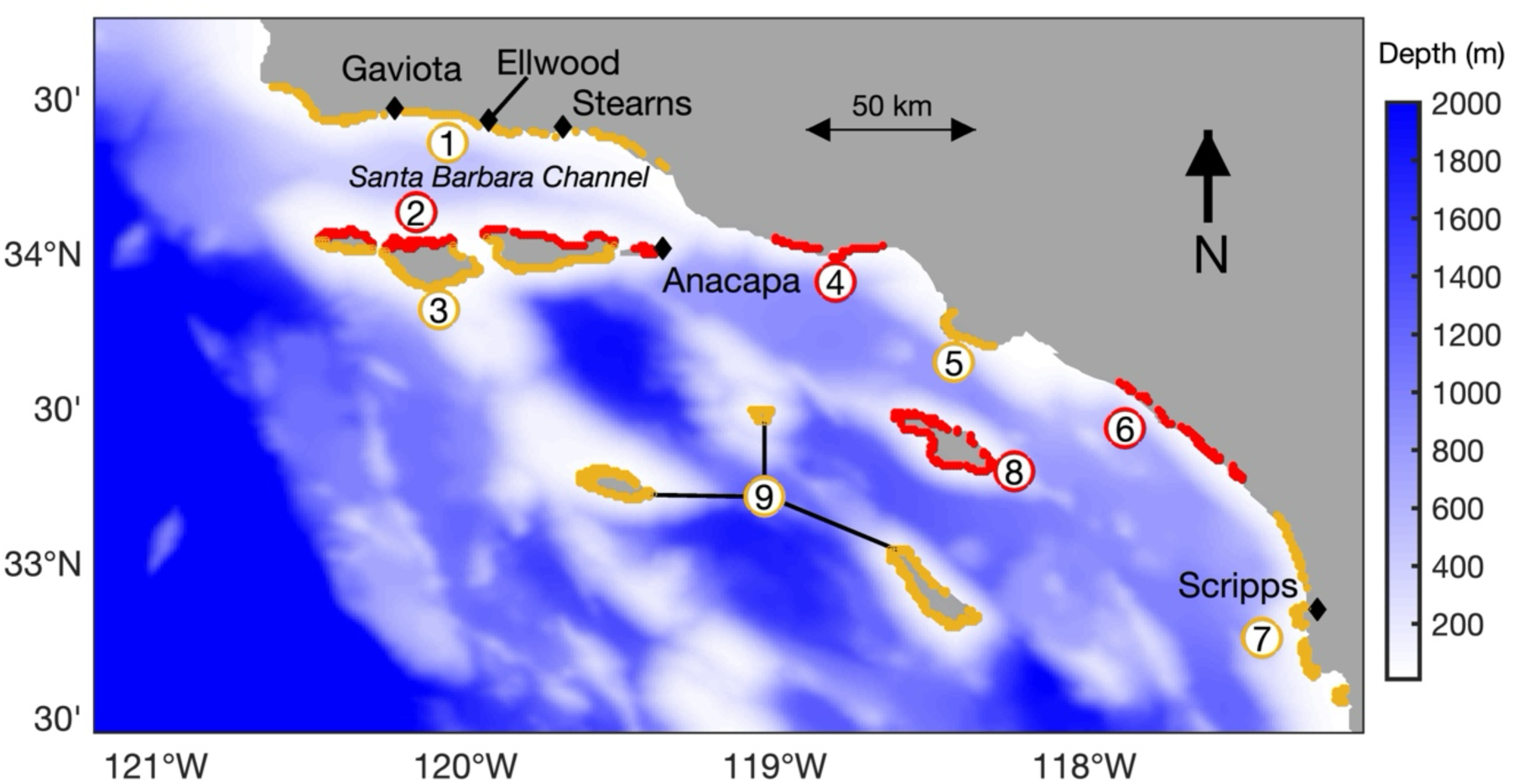
Southern California Bight: study area and model domain. Black diamonds are the 5 larval recruitment sites: Gaviota, Ellwood, Stearns, Anacapa, and Scripps. Red and yellow coastline locations are the nine source zones, which are identified by the black numbers: (1) mainland of the Santa Barbara Channel, (2) north side of the North Channel Islands, (3) south side of the North Channel Islands, (4-7) groups of contiguous source sites along the central and southern mainland of the Southern California Bight, (8) Catalina Island, and (9) three South Channel Islands

The SC Bight, the southern component of the California Current System, is a transition zone lying at the interface between two very different oceanographic regimes, the cold nutrient-rich waters of the south-flowing California Current and the warm nutrient-poor equatorial waters of the north-flowing Southern California Countercurrent (Lynn and Simpson 1987; Hickey 1993; Winant et al. 2003; Dong et al. 2009). The meeting of these two regimes in the SC Bight creates a year-round temperature gradient with colder water in the north and warmer water in the south and a seasonal chlorophyll gradient with high levels in the north and low levels in the south during the spring and early summer (Otero and Siegel 2004; Henderikx Freitas et al. 2017; Simons and Catlett 2023). As a result, the SC Bight is particularly sensitive to shifts in the California Current, which creates dramatic changes in ocean circulation, temperature, and primary productivity in the SC Bight and are often correlated with El Niño-Southern Oscillation (ENSO) events (Lynn and Bograd 2002; Dong et al. 2009). These features, in combination with an abundance of long-term ecological data, make the SC Bight an ideal system for studying the relationship between changing ocean conditions and larval recruitment.

In the California Current, changes in abundance of the purple sea urchins, *Strongylocentrotus purpuratus,* indirectly regulates the species assembly of nearshore rocky coastlines through voracious herbivory on kelp (Tegner and Dayton 1991; Filbee-Dexter and Scheibling 2014). A substantial body of research describes how predation and disease impact adult Strongylocentrotid urchin populations (Mann and Breen 1972; Estes and Duggins 1995; Lafferty 2004; Filbee-Dexter and Scheibling 2014; Burt et al. 2018) and documents climate-related booms and busts of purple urchins along the California Coast in recent years (Rogers-Bennett and Catton 2019; Okamoto et al. 2020; Rogers-Bennett and Okamoto 2020; Shanks et al. 2020). However, despite the widespread recognition of the importance of larval transport and recruitment in controlling population fluctuations in many marine species (Morgan 2001; Underwood and Keough 2001; Pineda et al. 2007; White et al. 2019), no studies have evaluated how changes in the myriad of ocean conditions influence larval recruitment of purple urchins in the California Current. The purple sea urchin (*S. purpuratus*) is used for this study because long-term, spatially replicated recruitment data is available at six locations in the SC Bight and prior work suggests that high temperatures and ENSO events in the SC Bight are linked to poor recruitment years (Okamoto et al. 2020) in part through changes in reproduction (Okamoto et al. Accepted).

The main goal of this study is to assess the impacts of changes in circulation, temperature and food supply on larval recruitment of purple sea urchins in the SC Bight. A biophysical model of the SC Bight, an ocean circulation model with particle tracking, is used to estimate larval recruitment at the recruitment sites where long-term urchin recruitment data has been collected (Fig. 1, Okamoto et al. 2020). The impact of changing circulation, temperature, and food supply on larval recruitment is quantified by deriving five indices that represent ocean conditions for each recruitment site. Two indices represent circulation, larval dispersal distance (LDD) and source zone strength (SZS). LDD is the distance a particle travels from a source site to a recruitment site. SZS is the relative probability that larvae would originate from nine coastal zones in the SC Bight (Fig. 1). Two indices represent temperature, larval temperature (T_L_) and source site temperature (T_S_). T_L_ is the temperature that larvae are exposed to as they are transported from the source sites to the recruitment sites. T_S_ is the temperature that adult urchins are exposed to at the spawning or larval source sites during reproduction. One index represents food availability (Chl), which is estimated using surface chlorophyll concentrations from satellite data along the particle tracks. The five ocean condition indices (OCI) are calculated for an 18-year modeling period from 1996-2013.

To accurately model larval transport and recruitment, the larval life history of the species must be realistically represented in the biophysical model (Paris et al. 2007; Cowen and Sponaugle 2009; Metaxas and Saunders 2009). However, empirical data on larval life history traits, such as vertical migration behavior, of many species is not available, and thus many studies do not include species-specific larval life history when modeling larval transport and connectivity (Swearer et al. 2019). To address this problem, we evaluate the sensitivity of the five OCI to variations in three larval life history parameters, plankton larval duration (PLD), larval behavior, and nearshore retention. Two types of PLD are simulated, a PLD of 42 days, which is an intermediate value from literature for purple sea urchins (Cameron and Schroeter 1980; Miller and Emlet 1999; Tegner 2001), and a temperature dependent PLD based on experimental studies. Two types of larval behavior are simulated passive drifting and diel vertical migration, which occurs when the larvae migrate to the surface at night and to depth during the day. For passive drifting, particles are moved by circulation only. Although evidence of vertical migration for echinoderms is very limited (Metaxas 2020; Doll et al. 2022), diel vertical migration has been widely observed for invertebrate larvae (Cohen and Forward 2009; Bandara et al. 2021) and is thus included in this study. With a 1 km^2^ horizontal scale, the ocean circulation model does not reproduce circulation patterns that are smaller than 1 km, including nearshore circulation which has been hypothesized to retain larvae (Morgan et al. 2009, 2018; Hagerty et al. 2018; Yamhure et al. 2021). Thus, a relationship is derived from the number of times a particle passes through a source site to represent nearshore retention.

Using all possible combinations of the three larval life history parameters, 12 simulations are conducted over the 18-year modeling period for each of five OCI, LDD, SZS, T_L_, T_S_ and Chl, determining how sensitive the OCI are to variations in the larval life history parameters. All OCI are found to be relatively insensitive to static versus dynamic PLDs, but showed varying levels of sensitivity to both nearshore retention and larval behavior. LDD and SZS are found to be more sensitive to nearshore retention and larval behavior than Chl, T_S_, and T_L_. To determine which OCI are the most likely drivers of larval recruitment in the SC Bight, all simulations of the OCI are statistically compared to a larval recruitment index derived from larval recruitment data (Okamoto et al. 2020). T_S_ is found to be the mostly likely driver to larval recruitment. LDD and SZS are found to potential drivers of larval recruitment if larvae are conducting DVM and/or influenced by nearshore retention, while T_L_ and Chl are unlikely to be drivers of larval recruitment.

## 2.0 Methods

### 2.1 Biophysical Model

To simulate larval transport, a 3D biophysical model consisting of two parts, an ocean circulation model (OCM) and a particle tracking model (PTM), was used. The OCM was a high-resolution Regional Ocean Modeling System (ROMS) applied to the SC Bight region (Dong and McWilliams 2007; Dong et al. 2009). The model domain, shown in Fig. 1, contained the SC Bight and had 1 km horizontal resolution with 40 vertical levels. The model was thoroughly validated for mesoscale circulation (Dong et al. 2009) and has successfully reproduced upwelling events in the SC Bight (Dong et al. 2011; Simons and Catlett 2023) and eddy circulation in the Santa Barbara Channel (SBC, Simons et al. 2015). The temperature from the OCM was validated using temperature data from 26 stations in the SC Bight (see Supplementary Information (SI)). Six-hour averaged 3D flow fields and temperature solutions produced by the OCM from 1996-2013 were used for this study.

Larval transport was simulated using the 3D PTM driven by 3D flow fields produced by the OCM (Mitarai et al. 2009; Simons et al. 2013). For this project, the particle tracking was run in reverse time, which was possible since the ROMS flow fields and temperature solutions were stored ofline. Reverse particle tracking was used to simulate the arrival of larvae at five recruitment sites in the SC Bight, Gaviota, Ellwood, Stearns, Anacapa, and Scripps, where larval recruitment has been recorded since 1993 (Fig. 1, Schroeter et al. 2017; Okamoto et al. 2020). The PTM has been used extensively for larval transport in the SC Bight (Mitarai et al. 2009; Simons et al. 2016; Nishimoto et al. 2019; Page 2019), and validated against observational data from drifter experiments (Ohlmann and Mitarai 2010).

Particles were released from 45 horizontal locations evenly spaced every square kilometer around each recruitment site. This distribution of horizontal release points was selected to capture the mesoscale circulation in the area surrounding the recruitment sites. At each of the 45 horizontal release points, particles were released vertically every meter from 1 to 30 meters below the surface, the maximum depth where purple sea urchins are typically found in high densities. By evenly distributing particles over the water column, the particle distribution reflected average vertical flow conditions as recommended for robustness in biophysical models (Simons et al. 2013). Particles were released every 6 hours from the 1,350 vertical and horizontal locations at each recruitment site for 1996-2013 and tracked for 8 weeks. The tracking time of 8 weeks was selected because it is the maximum possible plankton larval duration under idealized conditions (Munstermann et al. In Review). The location and temperature of every particle trajectory was recorded every 6 hours. The particle release frequency was selected to meet the particle number criteria for model robustness (Simons et al. 2013).

The particles arriving at the recruitment sites must originate from a source site in the study area (Fig. 1). Potential larval source sites in the SC Bight were identified by location of suitable spawning habitat for sea urchins. Suitable spawning habitat for sea urchins consists of rocky reef substrate located within areas that can hosts kelp or are algal dominated. Thus, the spatial distribution of larval source sites in the SC Bight was derived from satellite observations of giant kelp canopy (Bell et al. 2015). Using the mean distribution of giant kelp canopy from 1984-2022, suitable spawning habitat was found in 926 of the 1-km^2^ model grid cells, which were designated as potential larval source sites (Fig. 1). To represent the peak recruitment season for purple sea urchins, which runs from March to June in the SC Bight (Okamoto et al. 2020), particles were released every 6 hours from March 1^st^ to June 30^th^ for every year of the modeling period (1996-2013) at the release locations for each recruitment site. Based on the particle release locations and frequency, a total of 12 million particles were released from each recruitment site over the 18-year modeling period.

To quantify the horizontal dispersal of particles, the 3D particle tracks were transformed into 2D particle density maps (PDMs). Using the three-dimensional particle tracks for each recruitment month and site, 2D monthly PDMs were produced by compiling the horizontal locations of all particle tracks from the particles released over the month into a single dataset and then summing the number of particle locations within each model grid cell of 1 km^2^, creating a 2D particle density matrix the same size as the model grid. Example PDMs for Gaviota and Scripps sites are shown in Fig. 2(a-f).

**Figure 2:**
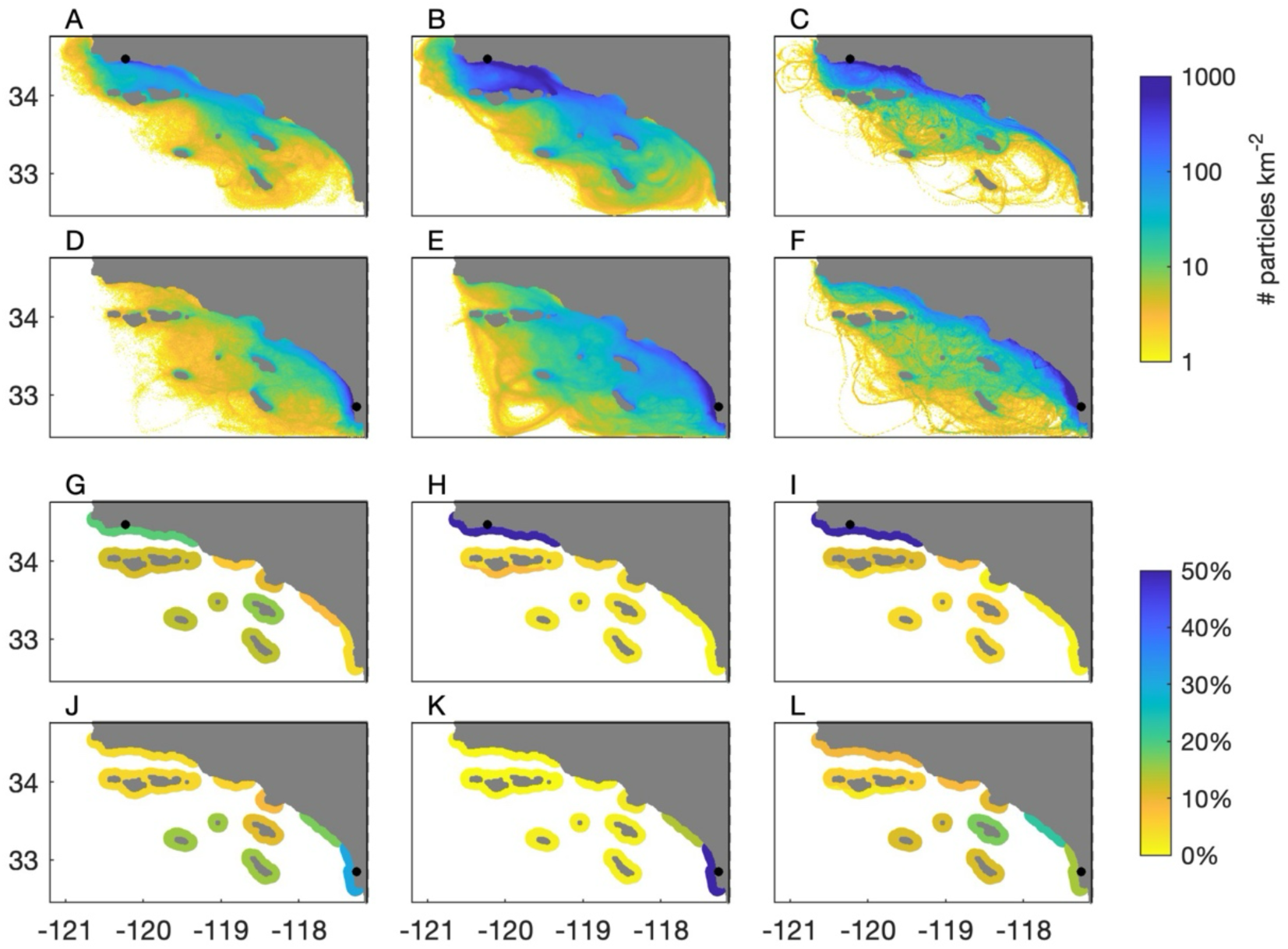
(A-F) Mean particle density maps (PDMs) and (G-L) mean source zone strength (SZS) for three following simulations with static PLDs: (1) passive drifting without nearshore retention for Gaviota (A,G) and for Scripps (D,J), (2) passive drifting with nearshore retention for Gaviota (b,h) and Scripps (E,K), and (3) DVM20 without nearshore retention for Gaviota (C,F) and for Scripps (F,L). PDMs and SZS have been averaged over all months of the modeling period (1996-2013). PDMs are plotted on a log scale.

### 2.2 Larval life history parameters

To accurately simulate the transport of purple sea urchin larvae in the SC Bight, larval life history parameters, which include larval behavior, planktonic larval duration (PLD), and nearshore retention, must be incorporated into the biophysical model. This section describes how each of these parameters was modeled.

Three types of larval behavior were simulated with the PTM, passive drifting, diel vertical migration to 20 m depth (DVM20), and diel vertical migration to 40 m depth (DVM40). To simulate passive drifting, particles were moved only by 3D currents. Diel vertical migration (DVM) occurs when larvae migrate to the surface at night and to depth during the day to feed and/or avoid predation and is hypothesized to be triggered by light availability (Cohen and Forward 2009; Bandara et al. 2021). As the vertical movement or distribution of sea urchin larvae has not been studied in the SC Bight, depths of 20 m and 40 m were selected to test DVM behavior. As the average depth of the euphotic zone in the SC Bight is ∼30 m (Brzezinski and Washburn 2011; Krause et al. 2013), depths of 20 m and 40 m were selected such that particles would migration within the euphotic zone and below euphotic zone respectively. DVM20 and DVM40 were modeled by particles remaining at near-surface depths (∼3 m) at night (18:00 hr to 06:00 hr) and at deeper depths of 20 m or 40 m during the day (06:00 hr to 18:00 hr). A basic random walk was used to vertically distribute the particles in a gaussian pattern around the set depths, near-surface, 20 m, and 40 m.

PLD is represented in the PTM by the length of time particles are tracked. Two types of PLD were used for this study. The first PLD was 6 weeks, which was an intermediate value based on both 4literature and experimental data (Cameron and Schroeter 1980; Miller and Emlet 1999; Tegner 2001) and was termed “static PLD”. The second PLD was a temperature dependent PLD based on experimental studies, which was termed “dynamic PLD”. To model the static PLD, the particle tracking time was set to six weeks or 42 days. To calculate the dynamic PLD, the following two-step process was used. First, using all particle tracks for one recruitment site and month of the modeling period, the mean particle temperature was calculated for daily tracking times of 1 to 56 days. Second, the dynamic PLD was determined by the intersection of two lines of temperature versus PLD, one line from model results and the other line from the experimental data (example shown Fig. S1). Experimental data indicates PLD as a function of temperature does not vary with food availability (Munstermann et al. In Review).This process was repeated for the three larval behaviors, five recruitment sites, and four months of the recruitment period for the 1996-2013 modeling period. The dynamic PLDs ranged from 26 to 56 days with a mean of 36 days and are displayed Fig. S2.

To quantify the effect of nearshore circulation on larval retention, two contrasting simulations, one without nearshore retention and one with nearshore retention, were produced. The simulation without nearshore retention assumed that particles travelled continuously from a source site to a recruitment site during their PLD and were not retained in a source site even if they passed through a source site during their PLD. As no information exists on the probability of larvae being retained along the shoreline, the simulations with nearshore retention assumed that the number of particles released during a recruitment month that passed through a source site at any time during their PLD was directly proportional to the probability of particles being retained within the source site. Thus, the simulations with nearshore retention contained all particle tracks that passed through a source site at any time during their PLD. If a particle track did not originate from a source site at the beginning of their PLD, the particle track was truncated at the first source site it entered, and the particle was then assumed to be retained within this source site for the portion of the PLD that it was not being transported to the recruitment site. Consequently, the PDMs without nearshore retention were created using only the particle tracks that start at a source site at the beginning of the their PLD, while the PDMs with nearshore retention were created using particle tracks that start a source site anytime during their PLD. This means that PDMs with nearshore retention include many particle tracks that are shorter than the PLD and assume that these particles are retained within the source site at the beginning of their PLD until they are transported to the recruitment site.

### 2.3 Ocean condition indices

This section describes how the five ocean condition indices (OCI), larval dispersal distance, source zone strength, larval food supply, larval temperature, and temperature at the source sites, were derived from particle tracking results. All OCI are calculated on a monthly timescale for the modeling period of 1996-2013. Larval dispersal distance, source zone strength, larval food supply, and larval temperature are calculated for the recruitment period of March to June. Temperature at the source sites is calculated for the reproductive period of September to November. For the larval life history parameters described in the previous section, only nearshore retention changed the particle tracking times to less than the PLD. This change is particle tracking times affected how source zone strength and larval temperature were calculated with and without nearshore retention and is explained in this section.

#### Larval dispersal distance (LDD)

To compare larval dispersal between simulations, a monthly LDD for each recruitment month and site was produced. The LDD was derived by calculating the distance between the source site and recruitment site for each particle and then averaging this distance for all particles tracked over the month. Thus, a monthly time-series of LDD for the recruitment period was produced for each recruitment site in units of kilometers. To derive a mean LDD for the entire 18-year study period, the LDD was averaged over all months and years of the study period and is termed “mean LDD”.

#### Source zone strength (SZS)

When using a reverse particle tracking method, source site strength is defined as the probability that a larva arriving at a recruitment site originated from a specified source site. For each recruitment sites, source site strength for the recruitment period was derived by summing the number of particles that originated from or passed through a source site for one month of the modeling period. Thus, the number of particles at each of the 926 source sites represents the relative probability or strength of the source sites. Without nearshore retention, source site strength was calculated using only the particles that originated at a source site at the beginning of the PLD and by counting each particle one time at its source site. With nearshore retention, source site strength was calculated using particles that originated at a source site anytime during the PLD and by counting the particle every time its track passed through a source site. Thus, the method for implementing nearshore retention resulted in particles being counted multiple times for different source sites. The source sites were then geographically grouped into nine source zones (Fig. 1). The nine source zones consisted of (1) the mainland of the SBC, (2) the north side of the North Channel Islands, (3) the south side of the North Channel Islands, (4-7) groups of contiguous source sites along the central and southern mainland of the SC Bight, (8) Catalina Island, and (9) three South Channel Islands. For each recruitment site and month, the percentage of particles in each zone was calculated, producing a monthly time-series of source zone strength (SZS) for each zone and recruitment site over the recruitment period.

#### Larval food availability (Chl)

To estimate food availability during larval transport, surface chlorophyll was used as a proxy for phytoplankton, which is what urchin larvae use for food (Strathmann 1975). Surface chlorophyll for the SC Bight was calculated from satellite-derived 5-day composites of surface chlorophyll-a concentrations (Kahru et al. 2012, 2015, https://spg-satdata.ucsd.edu) and was available from 1998 to 2013, 16 years of 18-year modeling period. As the satellite data is two-dimensional and does not represent the chlorophyll in the sub-surface, we recognize that this is a limitation of our analysis. However, surface chlorophyll has been shown to reflect the overall seasonal and interannual changes in chlorophyll in the SC Bight (Otero and Siegel 2004; Henderikx Freitas et al. 2017; Simons and Catlett 2023) and can provide some insight into the correlation between food availability and larval recruitment. In addition, satellite data was the only source of chlorophyll data that provided the spatial and temporal coverage needed to use with the particle tracking model results.

The 5-day composites of surface chlorophyll were averaged over each recruitment month of March-June to produce monthly surface chlorophyll of the SC Bight from 1998-2013. For each recruitment month and site, particle exposure to surface chlorophyll was calculated by first interpolating the model grid onto the monthly surface chlorophyll and then multiplying the surface chlorophyll in each model grid cell by the corresponding fraction of particles in the grid cell derived from the PDMs. Using this method, monthly time-series of larval exposure to chlorophyll (Chl) for each recruitment site was derived in units of mg m^-3^.

#### Larval temperature (T_L_)

Larval temperature (T_L_) represents the temperature that larvae are exposed to during their PLD. T_L_ was derived from two sources of temperature produced by the OCM for the recruitment period. The first source was the particle temperature, which was recorded every 6 hours along the particle track. The second source was the temperature at the source sites, which was calculated every six hours at 5 m depth in the model grid cell where the source site was located. A depth of 5 m was selected because it is a common depth of sea urchins in the SC Bight (Okamoto, D., personal observation). For simulations without nearshore retention, only particle temperature was used to calculate T_L_ as particles were tracked for the entire duration of the PLD. In this case, T_L_ was calculated by first averaging the particle temperature for each particle track, producing a mean particle temperature, and then averaging the mean particle temperatures for all the particles from one recruitment site and one month of the modeling period, producing a monthly time-series of T_L_ for each recruitment site in units of °C for the recruitment period. For simulations with nearshore retention, both particle temperature and temperature at the source sites were used to calculate T_L_ as many particles were retained at a source site for part of the PLD before being transported to a recruitment site. For the particles that were retained at a source site, the mean particle temperature was calculated by averaging the temperature at the source site for the time that the particle was retained there and the particle temperature for the time that the particle was tracked from the source site to the recruitment site. For the particles that were not retained at a source site, the mean particle temperature for particles was calculated the same way as the simulations without nearshore retention. Using both particles that were and were not retained at a source site, T_L_ for simulations with nearshore retention was derived by averaging the mean particle temperatures from one recruitment site and one month of the modeling period, producing a monthly time-series of T_L_ for each recruitment site in units of °C for the recruitment period.

#### Source site temperature (T_S_)

The reproductive period for purple urchins in the SC Bight occurs from September to November. To assess the impact of temperature on adult urchins during the reproductive period, temperature at the source sites (T_S_), the location of the adult urchins, was needed. T_S_ was calculated using the following four-step method. First, the monthly mean temperature at 5 m below the surface was calculated from the OCM at each of 926 source sites for the reproductive period from 1996-2013. Second, the monthly source site strength for the each of the 926 source sites was averaged over the recruitment period (March-June) for each year, producing yearly source site strength for each of the 18 years of the modeling period and each source site. Third, the yearly source site strength for each site was then divided by the sum of the yearly source site strength for all source sites, producing a fraction for each source site and year. As reproduction occurs prior to recruitment, the T_S_ during the reproductive period was calculated from the source site strength for the following year and thus, limited to a 17-year modeling period of 1996-2012. Fourth, using the fractions derived from source site strength and temperature at the source sites, a weighted mean temperature or T_S_ was calculated for each recruitment site and month of the reproductive period, producing a 17-year time-series of monthly T_S_ for each recruitment site in units of °C.

### 2.4 Sensitivity study of larval life history parameters

Sensitivity of the five OCI, LDD, SZS, Chl, T_L_, and T_S_, to variations in three larval life history parameters of PLD, larval behavior, and nearshore retention was tested. PLD had two options, static and dynamic. Behavior had three options, passive drifting, DVM20, and DVM40. Nearshore retention had two options, with and without nearshore retention. For each OCI, 12 simulations were created using all possible combinations of the three parameters and are listed in Table 1. Simulation 1 with passive drifting behavior, no nearshore retention, and a static PLD was designated as the baseline simulation as it represents how larval transport is simulated for many other biophysical modeling studies (Swearer et al 2019).

**Table 1:** Simulations created for each of the five ocean condition indices (OCI), LDD, SZS, Chl, T_L_, and T_S_, with three larval life history parameters of larval behavior, nearshore retention, and PLD.

| Simulation Number | Larval Life History Parameters |  |  |
| --- | --- | --- | --- |
|  | Larval Behavior | Nearshore Retention | PLD |
| 1 | Passive Drifting | No | Static |
| 2 | Passive Drifting | No | Dynamic |
| 3 | Passive Drifting | Yes | Static |
| 4 | Passive Drifting | Yes | Dynamic |
| 5 | DVM20 | No | Static |
| 6 | DVM20 | No | Dynamic |
| 7 | DVM20 | Yes | Static |
| 8 | DVM20 | Yes | Dynamic |
| 9 | DVM40 | No | Static |
| 10 | DVM40 | No | Dynamic |
| 11 | DVM40 | Yes | Static |
| 12 | DVM40 | Yes | Dynamic |

For each ocean condition index, simulations were compared using three statical measurements, root-mean-square deviation (RMSD), Pearson’s correlation coefficient (*r*), and mean error (ME). The RMSD was used to assess the absolute magnitude of deviation between the simulations. The *r* was used to assess the linear correlation between the simulations. ME was used to quantify the increase or decrease in OCI between the simulations. To calculate RMSD, *r*, and ME for two simulations, the OCI must be in the form of a 1D time-series, which is the case for LDD, T_L_, Chl, and T_S_. However, the SZS has 9 1D time-series, with each time-series corresponding to one zone. In order to calculate RMSD and *r* for two SZS simulations, the 9 SZS time-series for each simulation were arranged into a 2D matrix with zone versus time and then linearized into a 1D vector.

To test the sensitivity of each parameter, RMSD, *r*, and ME were calculated for two simulations where one parameter varied and the other two parameters were held constant. The two simulations are termed a ‘simulation pair’. For PLD, 6 simulations pairs were used where one simulation had a static PLD and the other had a dynamic PLD while larval behavior and nearshore retention were the same for both simulations. The 6 simulation pairs for PLD were: (1) simulations 1 and 2, (2) simulations 3 and 4, (3) simulations 5 and 6, (4) simulations 7 and 8, (5) simulations 9 and 10, and (6) simulations 11 and 12. For nearshore retention, 6 simulation pairs were used where one simulation included nearshore retention and the other did not while larval behavior and PLD were the same for both simulations. The 6 simulations pairs for nearshore retention were: (1) simulations 1 and 3, (2) simulations 2 and 4, (3) simulations 5 and 7, (4) simulations 6 and 8, (5) simulations 9 and 11, and (6) simulations 10 and 12. For larval behavior, 12 simulations pairs were used where one simulation had passive drifting and the other had DVM20 or DVM40 or where one simulation had DVM20 and the other had DVM40 while PLD and nearshore retention was the same for both pairs. The 12 simulation pairs for larval behavior were: (1) simulations 1 and 5, (2) simulations 2 and 6, (3) simulations 3 and 7, (4) simulations 4 and 8, (5) simulations 1 and 9, (6) simulations 2 and 10, (7) simulations 3 and 11, (8) simulations 4 and 12, (9) simulations 5 and 9, (10) simulations 6 and 10, (11) simulations 7 and 11, and (12) simulations 8 and 12.

### 2.5 Correlation of ocean condition indices to larval recruitment index

To determine which ocean condition index was most likely driving larval recruitment in the SC Bight, the OCI were compared to an annual larval recruitment index that was derived from larval recruitment data. Recruitment indexes in the form of standardized larval settlement anomaly were available for the 5 recruitment sites including an additional site, Ocean Beach, which was located close to the Scripps recruitment site (Okamoto et al. 2020). A larval recruitment index was calculated for the northern SC Bight by averaging the indices from the four SBC recruitment sites, Gaviota, Ellwood, Stearns, and Anacapa, and for southern SC Bight by averaging the indexes from the recruitment sites for Scripps and Ocean Beach sites (Table S1). Northern and southern SC Bight recruitment indexes were developed due to the synchronous patterns of recruitment at the northern versus the southern recruitment sites observed by Okamoto et al. (2020), indicating that forces driving recruitment are on a regional versus a local scale. As the recruitment indices were on an annual time scale, the monthly time-series’ of the OCI were averaged over the recruitment period for LDD, SZS, Chl, and T_L_, and over reproductive period for T_S_ for each year to produce annual time-series’. The annual time-series of the OCI were then compared to the northern recruitment index for the Gaviota, Ellwood, Stearns, and Anacapa recruitment sites and the southern recruitment index for the Scripps recruitment site by calculating *r* for the 12 simulations. LDD, T_L_, Chl, and T_S_ were in the form 1D time-series and could be used directly to calculate *r*, but SZS was a 2D time-series and had to be converted into a 1D time-series to compare to the 1D larval recruitment index. To do this, 1D SZS for each simulation and site was calculated by summing SZS over all possible combination of one to three zones. For the 9 zones, 129 combinations of 1 to 3 zones were used, producing 129 1D SZS time-series for each simulation and site. Then *r* was calculated between the 129 1D SZS time-series and the larval recruitment index. To indicate that any of the five OCI may be driving recruitment, *r* would have to be of a similar high magnitude and sign across all recruitment sites for a single simulation.

## 3.0 Results

Our results are presented in two parts: first, the sensitivity of OCI to the larval life history parameters and second, the correlation of OCI to the larval recruitment index. The five OCI, which are LDD, SZS, Chl, T_L_, and T_S_, fall into three categories: ocean circulation represented by LDD and SZS, food supply represented by Chl, and temperature represented by T_L_ and T_S_. The larval life history parameters include PLD, nearshore retention, and behavior. PLD has two options, a static PLD of 42 days and a dynamic PLD, which is as a function of temperature. Nearshore retention has two options, included or not included in the simulation. Behavior has three options, passive drifting, DVM with daytime depth of 20m (DVM20), and DVM with a daytime depth of 40m (DVM40). Using all possible combinations of these larval life history parameters, 12 particle tracking simulations, shown in Table 1, are produced for each of the OCI and the recruitment sites over the 18-year modeling period of 1996-2013. Due to their location, the Gaviota, Ellwood, Stearns, and Anacapa sites are referred to collectively as the SBC sites, and the Gaviota, Ellwood, and Stearns sites are referred to collectively as the SBC mainland sites. Figures of the five OCI for the five recruitment sites and the 12 simulations are included in the SI (Fig. S3-S11).

### 3.1 Sensitivity of ocean condition indices to larval life history parameters

#### Larval dispersal distance

Larval dispersal distance (LDD) represents the mean distance that the larvae have traveled from the source sites to the recruitment sites and reflects changes in circulation. For the baseline simulation of passive drifting behavior, a static PLD, and no nearshore retention, LDD is shown for the recruitment period of March-June in Fig. 3(a,b) for Gaviota and Scripps. Gaviota and Scripps are shown because they are the recruitment sites that are farthest apart and represent the northern versus southern conditions in the SC Bight. For the baseline simulation, LDD is extremely variable with each recruitment site ranging an order of magnitude over the modeling period. Baseline LDD for the modeling period has a mean and standard deviation of 79.5 km and 51.6 km for Gaviota, 84.8 km and 39.6 km for Ellwood, 99.7 km and 30.8 km for Stearns, 71.1 km and 25.3 km for Anacapa, and 88.4 km and 47.8 km for Scripps.

**Figure 3:**
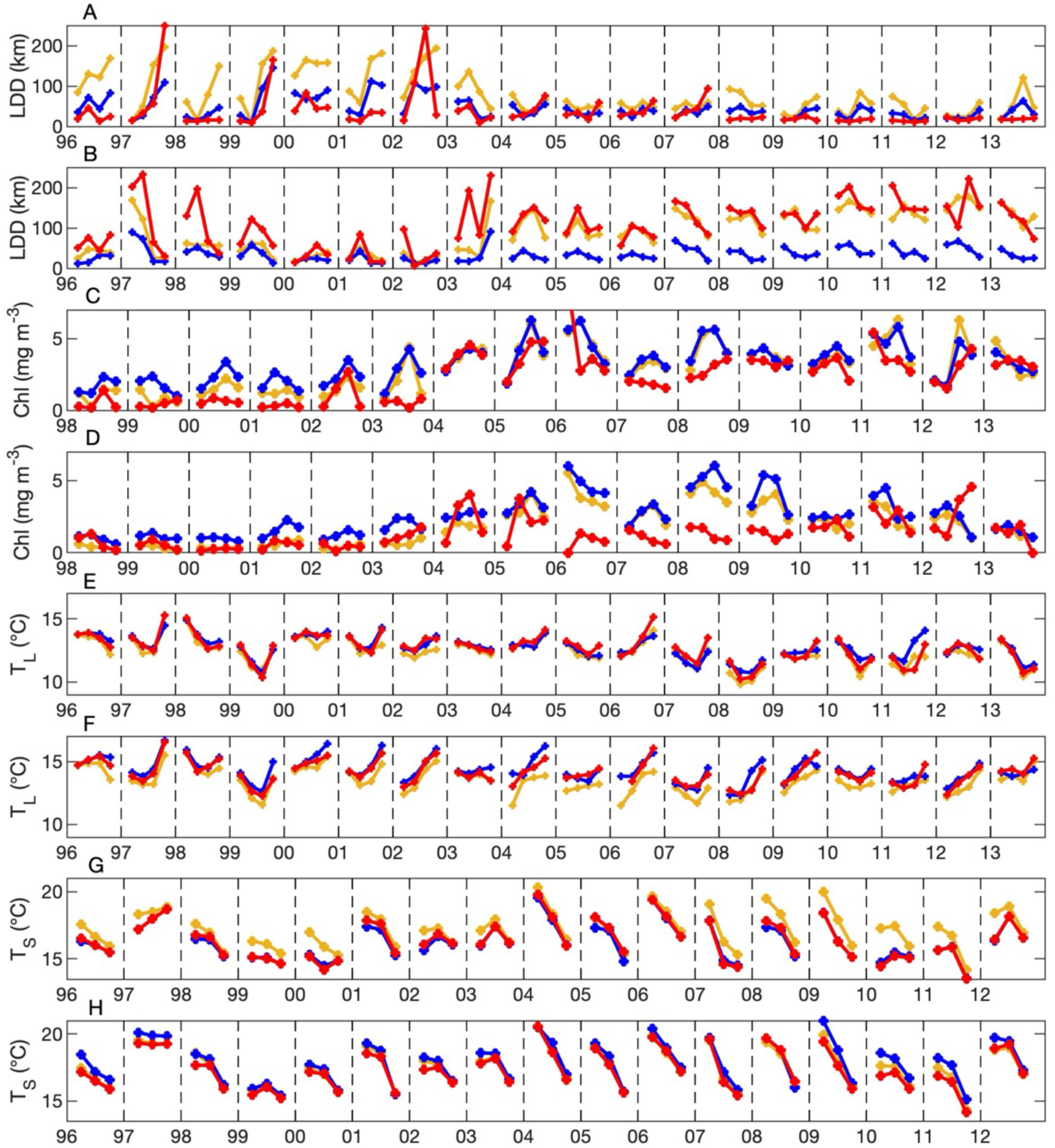
Ocean conditions for three simulations: (1) passive drifting behavior, no nearshore retention, and static PLD (yellow line), (2) passive drifting behavior, nearshore retention, and static PLD (blue line), and (3) DVM20 behavior, nearshore retention, and static PLD (red line). (A) LDD (km) for Gaviota, (B) LDD (km) for Scripps, (C) Chl (mg m^-3^) for Gaviota, (D) Chl (mg m^-3^) for Scripps, (E) T_L_ (°C) for Gaviota, (F) T_L_ (°C) for Scripps, (G) T_S_ (°C) for Gaviota, and (H) T_S_ (°C) for Scripps. LDD, Chl, and T_L_ are shown for the larval recruitment period of March-June. T_S_ is shown for the reproductive period of September-November.

##### Planktonic larval duration

To examine the sensitivity of LDD to PLD, the six simulations pairs with static versus dynamic PLD are compared and show very strong agreement for all recruitment sites. For all recruitment sites, RMSD ranges 1.8-16.9 km, *r* ranges from 0.95-0.99, and ME ranges from 0.01-10 km.

##### Nearshore retention

For nearshore retention, the sensitivity testing of LDD for all recruitment sites shows a strong linear correlation between all simulation pairs with and without nearshore retention for all sites, resulting in a *r* ranging from 0.73-0.96 with a mean of 0.88, but displays a large range of agreement in magnitude among the simulation pairs as a function of behavior and site location. For the SBC sites, the weakest agreement for LDD is observed for the two passive drifting simulation pairs. The two simulation pairs with passive drifting have RMSD ranging from 33.5-44.1 km for Gaviota, Ellwood, and Stearns and from 19.2-19.8 km for Anacapa. ME shows a decrease in LDD of 28.2-34.6 km for Gaviota, Ellwood, and Stearns and 14.0-14.5 km for Anacapa when nearshore retention is included in the simulation with passive drifting. The decrease in LDD for Gaviota when nearshore retention is included with passive drifting is shown in Fig. 3(a). For SBC sites, stronger agreement is observed for the four DVM20 and DVM40 simulation pairs with RMSD ranging from 11.4-25.6 km. ME for the four DVM20 and DVM40 simulations pairs shows a decrease in LDD of 4.6-10.9 km when nearshore retention is included, which is a smaller decrease in LDD than for the two passive drifting pairs. For Scripps, LDD shows similar magnitudes of agreement for all six simulation pairs with RMSD ranging from 55.7-67.0 km. ME for the four simulation pairs with passive drifting and DVM20 show a decrease in LDD of 44.9-54.0 km when nearshore retention is included, but the two simulation pairs with DVM40 show an increase in LDD of 19.2-21.8 km when nearshore retention is included. The decrease in LDD for Scripps when nearshore retention is included with passive drifting is shown in Fig. 3(b).

##### Larval behavior

For the SBC mainland sites, LDD shows different patterns of sensitivity in magnitude and linear correlation as functions of behavior and nearshore retention. For magnitude, LDD shows the weakest agreement for the four simulation pairs comparing passive drifting behavior to DVM20 or DVM40 without nearshore retention, medium agreement for the four simulation pairs comparing passive drifting behavior to DVM20 or DVM40 with nearshore retention, and the strongest agreement in LDD for the four simulation pairs comparing DVM20 to DVM40 with and without nearshore retention. The four simulation pairs comparing passive drifting behavior to DVM20 or DVM40 without nearshore retention have the largest RMSD ranging from 60.2-74.3 km. Fig. 3(a) shows LDD for Gaviota for simulations with passive drifting and DVM20 without nearshore retention. The four simulation pairs comparing passive drifting behavior to DVM20 or DVM40 with nearshore retention have smaller RMSD ranging from 27.9-42.9 km. The four simulation pairs comparing DVM20 to DVM40 with and without nearshore retention have the smallest RMSD ranging from 14.8-32.4 km. For the eight simulation pairs comparing passive drifting with DVM with and without nearshore retention, *r* ranges widely from 0.30 and 0.69, and the strength of correlation is not associated with larval behavior or nearshore retention. The four simulation pairs comparing DVM20 to DVM40 with and without nearshore retention have the highest *r* ranging from 0.71-0.91. For the eight simulations pairs with passive drifting and DVM, LDD has a ME ranging from 17.8-54.6 km, indicating that LDD is smaller for simulations with DVM than for simulations with passive drifting. The absolute ME between the four simulations pairs with DVM20 and DVM40 ranges from 4-12 km and does not show a consistent increase or decrease in LDD, indicating that LDD has similar magnitudes for DVM20 and DVM40 simulations.

For Anacapa, LDD shows similarly strong agreement for all simulations pairs with RMSD ranging from 20.1-28.7 km and *r* ranging from 0.56-0.75. The absolute ME for all simulations pairs is small, ranging from 0 to 15.4 km with a mean of 8.1 km. This small ME implies that there is no consistent increase or decrease in LDD associated with a specific behavior or nearshore retention.

For Scripps, LDD shows the largest range of agreement in magnitude and linear correlation for the simulation pairs of all the five recruitment sites. The four simulation pairs comparing passive drifting to DVM20 or DVM40 with nearshore retention have the weakest agreement with RMSD ranging from 115-150 km and *r* ranging from 0.29-0.42. The four simulation pairs comparing passive drifting to DVM20 or DVM40 without nearshore retention have stronger agreement with RMSD ranging from 55-79 km and *r* ranging from 0.49-0.63. Fig. 3(b) shows LDD for Scripps for simulations with passive drifting and DVM20 without nearshore retention. The four simulation pairs comparing DVM20 to DVM40 with and without nearshore retention have the strongest agreement with RMSD ranging from 27-40 km and *r* ranging from 0.79-0.88. The ME for Scripps shows an increase in LDD of 20-120 km for all simulation pairs, which means that LDD is larger for simulations with DVM20 or DVM40 than passive drifting and for simulations with DVM40 than DVM20 regardless if nearshore retention is present. The magnitude of ME for Scripps does not correlate with a specific behavior or nearshore retention.

#### Larval source zone strength

Larval source zone strength (SZS) is defined as the probability that larvae have originated from a specified zone and represents how ocean circulation drives spatial and temporal variation in SZS. The nine source zones are identified in the SC Bight (Fig. 1). The units for SZS are the percentage of particles per zone, which is directly proportional to SZS. SZS for the baseline simulation of passive drifting, no nearshore retention, and a static PLD is shown for the recruitment period of March-June in Fig. 4(a,d) for Gaviota and Scripps. For all recruitment sites, baseline SZS is extremely variable in time and space over the 9 zones. Baseline SZS for Gaviota is similar to Ellwood, Stearns, and Anacapa with RMSD ranging from 5-9% and *r* ranging from 0.72-0.90, but very different to Scripps with a RMSD of 18% and a *r* of 0.02. For the baseline simulation, the maximum SZS per month and zone has a mean and standard deviation of 36% and 18% for Gaviota, 34% and 14% for Ellwood, 34% and 13% for Stearns, 32% and 13% for 40% for Anacapa and 20% for Scripps. Maximum SZS occurs predominantly in Zones 1, 3, 8, and 9 for Gaviota, Ellwood, Stearns, and Anacapa and Zones 6, 7, and 9 for Scripps.

**Figure 4:**
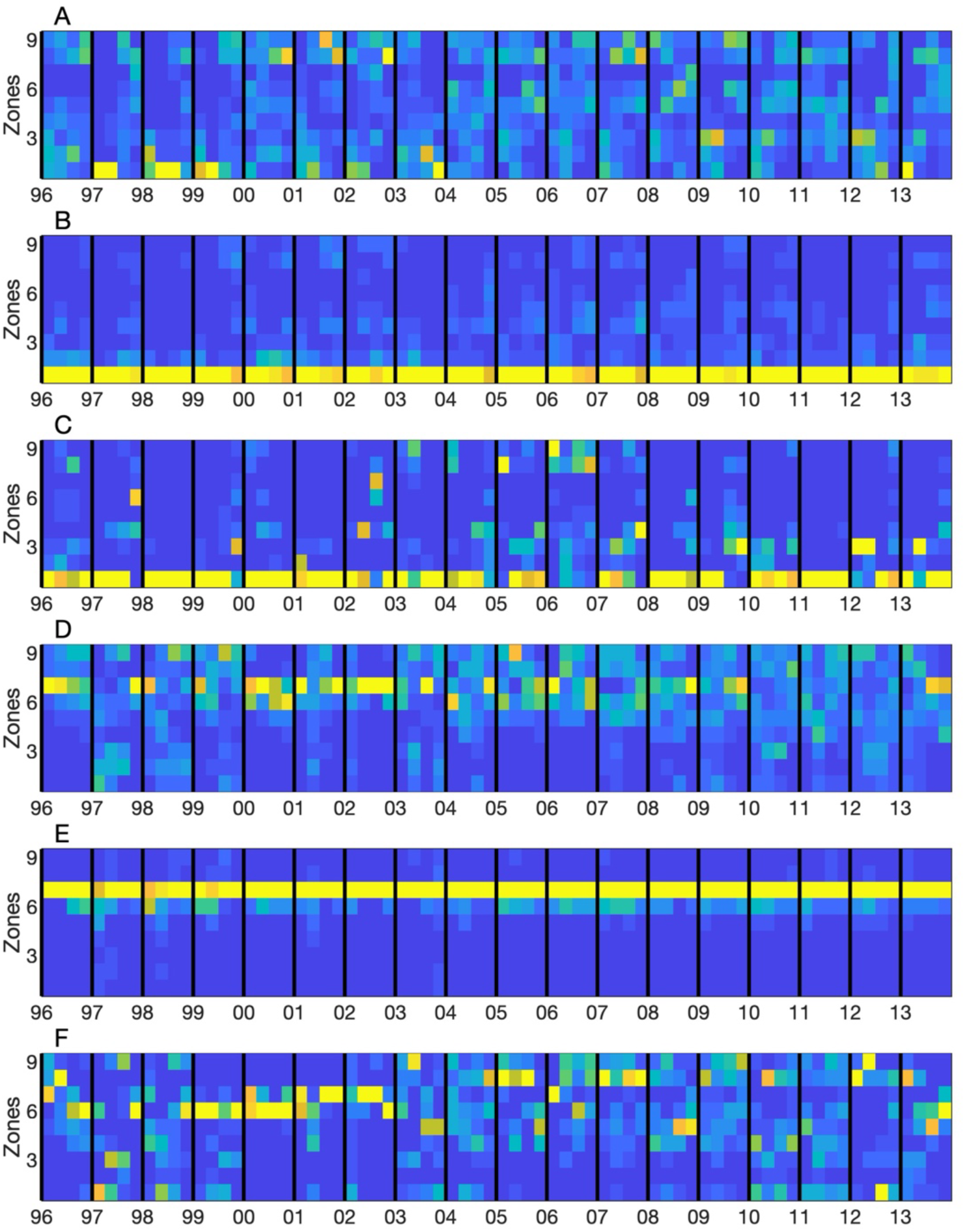
SZS for three simulations, (S1) passive drifting behavior, no nearshore retention, and static PLD, (S2) passive drifting behavior, nearshore retention, and static PLD, and (S3) DVM20 behavior, nearshore retention, and static PLD: (A) S1 for Gaviota, (B) S2 for Gaviota, (C) S3 for Gaviota, (D) S1 for Scripps, (E), S2 for Scripps, and (F) S3 for Scripps. SZS is shown for the recruitment period of March-June. Units for SZS is in percentage of particles. Zones 1-9 are shown in Figure 1.

##### Planktonic larval duration

The sensitivity testing of SZS to PLD shows strong agreement for all simulation pairs with a static versus a dynamic PLD for all recruitment sites. For all simulation pairs and recruitment sites, the RMSD ranges 1-9%, the *r* ranges from 0.88-1.0, and the ME ranges from 0-8%.

##### Nearshore retention

For the SBC mainland sites, SZS shows similar patterns of agreement in magnitude and linear correlation as function of behavior. SZS shows the weakest agreement for the two simulation pairs comparing passive drifting with and without nearshore retention, producing RMSD ranging from 19-21% and *r* ranging from 0.21-0.37. As shown in Fig. 2(g,h) and Fig. 4(a,b) for Gaviota, the addition of nearshore retention to passive drifting produces in the highest SZS occurring in the Zone 1, where the recruitment sites are located. For the two passive drifting simulation pairs, the ME in Zone 1 shows SZS increasing by 46-53% when nearshore retention is present. SZS shows the strongest agreement for the four simulation pairs comparing DVM20 or DVM40 with and without nearshore retention, producing RMSD ranging from 12-15% and *r* ranging from 0.85-0.91. For the four DVM simulation pairs, the ME in Zone 1 shows SZS increasing by only 18-26% when nearshore retention is present.

For Anacapa, SZS shows similar agreement in magnitude for all simulation pairs with a RMSD ranging from 11-14%. The SZS for Anacapa shows a weaker linear correlation for the two passive drifting simulation pairs of *r* from 0.43-0.44 than the four DVM20 and DVM40 simulation pairs of *r* from 0.73-0.81. ME for Zone 2, where Anacapa is located, displays an increase in SZS by 28-29% when nearshore retention is added to the passive drifting simulations. ME for Zone 2 shows a smaller increase in SZS of 11-17% when nearshore retention is added the DVM simulations.

For Scripps, the magnitude of SZS is very similar for all simulations pairs with RMSD ranging from 18-20%. The linear correlation of SZS is slightly higher for the passive drifting simulation pairs, ranging from 0.59-0.69, than the DVM simulation pairs, ranging from 0.43-0.49. For passive drifting and DVM, the SZS for Zone 7 increases when nearshore retention is present as shown in Fig. 2(j,k) and Fig. 4(d,e). ME for Zone 7 shows a similar increase in SZS of 39-48% when nearshore retention is added the passive drifting and DVM simulations.

##### Larval behavior

For the SBC mainland sites, the sensitivity testing of SZS to behavior shows the weakest agreement for the four simulation pairs comparing passive drifting to DVM20 or DVM40 without nearshore retention, producing RMSD ranging from 22-26% and *r* ranging from 0.10-0.36. The passive drifting simulations without nearshore retention produce similarly low levels of SZS across all zones (Fig. 2(g) and Fig. 4(a) for Gaviota), while the DVM20 and DVM40 simulations without nearshore retention produce the highest SZS in Zone 1, the zone where the recruitment site is located (Fig. 2(i) and Fig. 4(c) for Gaviota). The ME in Zone 1 shows that SZS for DVM without nearshore retention is higher by 44-55% than SZS for passive drifting without nearshore retention. The sensitivity testing of SZS displays the strongest agreement between the eight simulation pairs comparing DVM20 to DVM40 with and without nearshore retention and passive drifting to DVM20 or DVM40 with nearshore retention, producing a RMSD ranging from 4-13% and a *r* ranging from 0.90-99. The ME in Zone 1 shows that SZS from DVM simulations with nearshore retention is higher by 17-19% than SZS from passive drifting simulation with nearshore retention and that SZS from DVM40 simulations is higher by 1-8% than SZS from DVM20 simulations with and without nearshore retention.

For Anacapa, testing the sensitivity of SZS to behavior shows the strongest agreement for the four simulations pairs comparing DVM20 to DVM40 with and without nearshore retention, producing RMSD ranging from 8-12% and *r* ranging from 0.80-0.88. The ME in Zone 2 shows that SZS from DVM40 simulations is higher by only 2-7% than SZS from DVM20 simulations. For the eight simulations pairs comparing passive drifting to DVM20 or DVM40 with and without nearshore retention, SZS for Anacapa shows strong agreement in magnitude with RMSD ranging from 14-18%, but weak linear correlation with *r* ranging from 0.35-0.60. The ME in Zone 2 shows that SZS from DVM simulations is smaller by 8-27% than SZS from passive drifting simulations. This pattern is the opposite of what is observed for the SBC mainland sites where SZS in Zone 1, the location of the recruitment sites, is higher for DVM than passive drifting. For Anacapa, the ME for Zones 1 and 3 shows that SZS from DVM simulations is higher by 17-25% and 6-13% respectively than SZS from passive drifting simulations.

For Scripps, the sensitivity testing of SZS shows the weakest agreement for the four simulations pairs comparing passive drifting to DVM20 or DVM40 without nearshore retention, producing RMSD ranging from 16-18% and *r* ranging from 0.43-0.50. For these four simulation pairs, the ME is largest for Zone 7 and shows SZS decreasing by a 16-22% when DVM is present versus passive drifting without nearshore retention. SZS shows strong agreement of similar magnitude for the eight simulations pairs comparing passive drifting to DVM20 or DVM40 with nearshore retention and DVM20 to DVM40 with or without nearshore retention, producing RMSD ranging from 4-13% and *r* ranging from 0.78-0.98. The ME in Zone 7 shows that SZS is higher for DVM simulations with nearshore retention by 22-27% for passive drifting simulations with nearshore retention and that SZS for DVM40 simulations is only 1-2% than DVM20 simulations.

#### Larval food availability

Larval food is represented by surface chlorophyll (Chl), which was available from 1998-2013, 16 years of the modeling period. For the five recruitment sites, Chl for the baseline simulation of passive drifting behavior, a static PLD, and no nearshore retention, is shown in Fig. 3(c,d) for the recruitment period of March-June for Gaviota and Scripps. Baseline Chl is most similar between the Gaviota site and the Ellwood and Stearns sites with a RMSD of 0.70 mg m^-3^ and 0.75 mg m^-3^ and a *r* of 0.94 and 0.90 respectively and least similar between the Gaviota site and the Anacapa and Scripps sites with a RMSD of 1.24 mg m^-3^ and 1.57 mg m^-3^ and a *r* of 0.88 and 0.74 respectively. Averaged over the modeling period, mean baseline Chl is 2.95 mg m^-3^ for Gaviota, 2.65 mg m^-3^ for Ellwood, 2.74 mg m^-3^ for Stearns, 2.06 mg m^-3^ for Anacapa, and 1.84 mg m^-3^ for Scripps. The similar mean baseline Chl between the SBC mainland sites of Gaviota, Ellwood, and Stearns and the decreasing mean baseline Chl from Gaviota to Anacapa to Scripps reflects the mean north-south gradient of Chl in the SC Bight (Fig. 5(a)).

**Figure 5:**
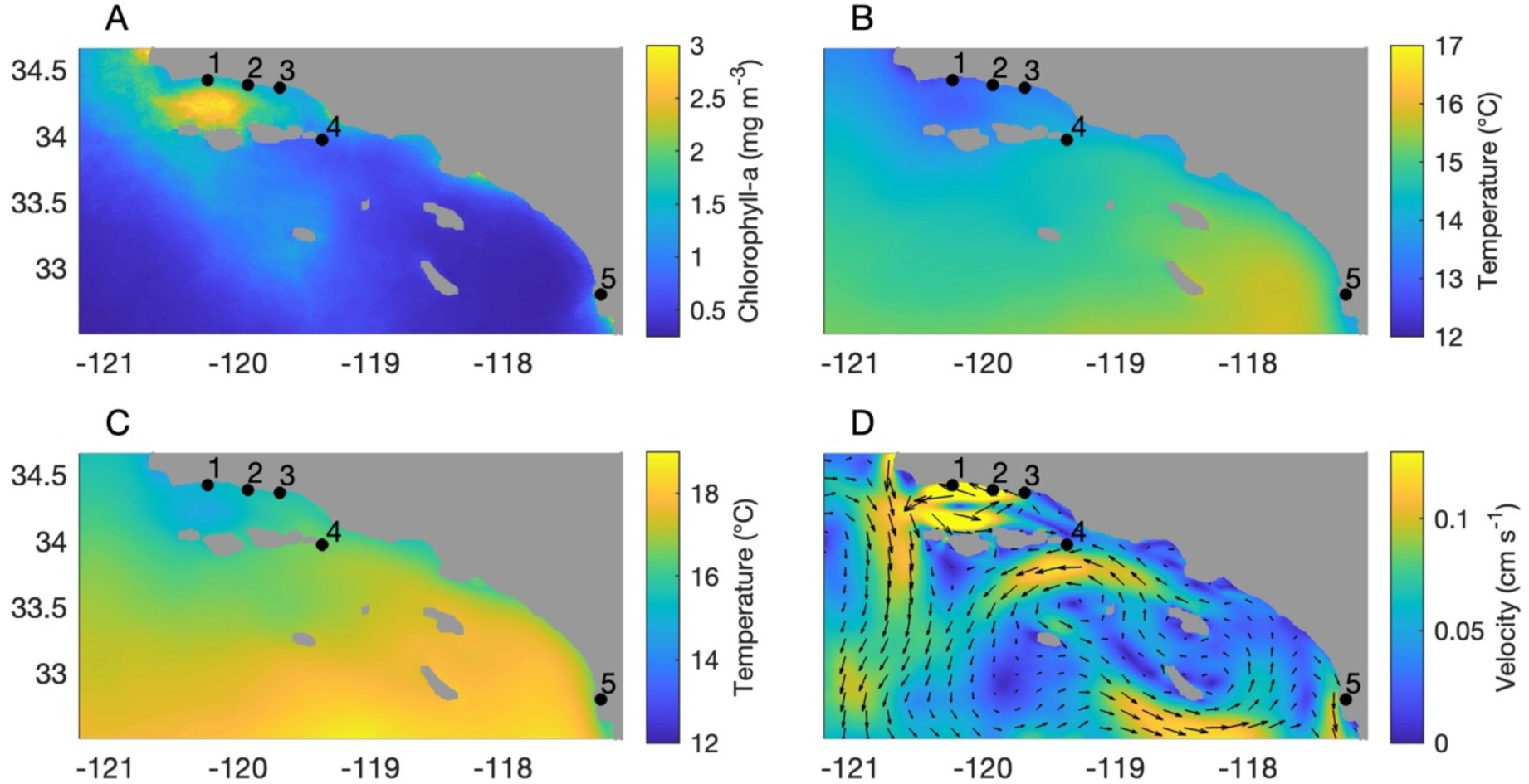
(A) Mean surface chlorophyll-a (mg m^-3^) from satellite data for the recruitment period, (B) mean temperature (°C) averaged over top 50 m of water column for the recruitment period, (C) mean temperature (°C) averaged over top 50 m of water column for the reproductive period, and (D) mean velocity (cm s^-1^) averaged over the top 50 m of the water column for the recruitment period. The solid black circles represent the recruitment sites: (1) Gaviota, (2) Ellwood, (3) Stearns, (4) Anacapa, and (5) Scripps. All data is averaged over the modeling period of 1996-2013. The recruitment period is from March to June. The reproductive period is from September to November.

##### Planktonic larval duration

The sensitivity testing of Chl to PLD shows strong agreement between all simulation pairs with a static PLD versus a dynamic PLD. For the six simulation pairs tested, RMSD ranges 0.10-0.37 mg m^-3^, *r* ranges from 0.98-1.0, and ME ranges from 0.01-0.20 mg m^-3^.

##### Nearshore retention

For the six simulation pairs with and without nearshore retention, the sensitivity testing of Chl for all recruitment sites shows strong linear correlations with *r* ranging from 0.91-1.0, but displays a range of agreement in magnitude among the simulation pairs as a function of behavior. For all recruitment sites, Chl shows a slightly weaker agreement in magnitude for the two passive drifting simulations pairs with RMSD ranging from 0.49-0.64 mg m^-3^ for the SBC sites and 0.76-0.81 mg m^-3^ for Scripps (Fig. 3(c,d)) than the four DVM20 and DVM40 simulation pairs with RMSD ranging from 0.29-0.46 mg m^-3^ for the SBC sites and 0.44-0.66 mg m^-3^ for Scripps. For all sites, the ME for Chl shows that the passive drifting simulations with nearshore retention are higher by 0.22-0.68 mg m^-3^ than the passive drifting simulations without nearshore retention. For the SBC sites, Chl is similar in magnitude for DVM simulations with and without nearshore retention with ME ranging from 0.02-0.09 mg m^-3^. For Scripps, the ME shows that Chl from the DVM simulations with nearshore retention are higher by 0.19-0.43 mg m^-3^ than Chl from the DVM simulations without nearshore retention.

##### Larval behavior

Chl shows strong linear correlations for the 12 larval behavior simulation pairs for all recruitment sites with *r* ranging from 0.80-0.99, but differences in magnitude as a function of behavior, nearshore retention, and site location. For the SBC sites, Chl shows the weakest agreement for the four simulation pairs comparing passive drifting to DVM20 or DVM40 without nearshore retention, producing RMSD ranging from 1.23-1.62 mg m^-3^. For these four simulation pairs, the ME shows that Chl for the DVM simulations is greater by 0.32-0.85 mg m^-3^ than the passive drifting simulations. For the SBC sites, Chl shows the slightly stronger agreement for the four simulation pairs comparing passive drifting to DVM20 or DVM40 with nearshore retention, producing RMSD ranging from 0.67-1.45 mg m^-3^. For these four simulation pairs, the ME shows that Chl for the DVM simulations is greater by 0.03-0.52 mg m^-3^ than the passive drifting simulations. For the SBC sites, Chl shows the strongest agreement for the four simulation pairs comparing DVM20 and DVM40 with RMSD ranging from 0.04-0.38 mg m^-3^ and ME from 0.01-0.12 mg m^-3^. For Scripps, Chl shows a range of weak to strong agreement for the 12 simulations pairs, but the magnitude of agreement is not correlated with larval behaviors or nearshore retention. For Scripps, Chl shows a RMSD ranging from 0.27-0.74 mg m^-3^ and ME ranging from 0.10-0.41 mg m^-3^.

#### Larval temperature

Larval temperature (T_L_) is defined as the temperature that larvae are exposed to during their PLD. T_L_ for the baseline simulation of passive drifting behavior, a static PLD, and no nearshore retention is shown in Fig. 3(e,f) for the recruitment period of March-June for Gaviota and Scripps. For the baseline simulation, T_L_ for all months and recruitment sites is below 16°C and ranges from 9.7-15.6°C. For all recruitment sites, the yearly change in baseline T_L_ over the recruitment period averages 1.5°C with a maximum of 2.8°C and does not show a consistent pattern of warming or cooling. Baseline T_L_ is very similar between mainland SBC sites of Gaviota, Ellwood, and Stearns. Comparing baseline T_L_ for the Gaviota site to the Ellwood and Stearns sites, RMSD and *r* range from 0.28-0.45°C and 0.92-0.96 respectively. Greater differences in baseline T_L_ are observed between Gaviota and Anacapa with a RMSD of 0.75°C and a *r* of 0.86 and between Gaviota and Scripps with a RMSD of 1.4°C and a *r* of 0.59. Averaging over the modeling period, mean baseline T_L_ is 12.4°C, 12.9°C, and 13.5°C for Gaviota, Anacapa, and Scripps respectively, reflecting the mean north-south temperature gradient in the SC Bight (Fig. 5(b)).

##### Planktonic larval duration

The sensitivity testing of T_L_ to PLD shows strong agreement between the six simulation pairs comparing static versus dynamic PLD. For the six simulation pairs and all recruitment sites, the RMSD ranges 0.13-0.36°C, the *r* ranges from 0.96-0.99, and ME ranges from 0.00-0.18°C.

##### Nearshore retention

The sensitivity testing of T_L_ to nearshore retention shows strong linear correlations for all simulation pairs and recruitment sites with *r* ranging from 0.83-0.98. However, differences in magnitude of T_L_ are observed with the weakest agreement for the simulation pairs with passive drifting (Fig. 3(e,f)) and the strongest agreement for the simulation pairs with DVM20 or DVM40 for all recruitment sites. T_L_ for the two simulation pairs with passive drifting have a RMSD ranging from 0.54-0.80°C for the SBC sites and a RMSD ranging from 0.87-1.0°C for Scripps. In contrast, the four simulation pairs with DVM20 or DVM40 have smaller RMSD ranging from 0.21-0.44°C for the SBC sites and a RMSD ranging from 0.55-0.82°C for Scripps. ME for T_L_ did not show consistent patterns as a function of nearshore retention or recruitment site.

##### Larval behavior

The sensitivity testing of T_L_ to behavior shows strong linear correlations for all simulation pairs and recruitment sites with *r* ranging from 0.85-0.99, but varies in magnitude of agreement as a function of behavior and site location. For the SBC mainland sites, T_L_ shows the weakest agreement for the four simulation pairs comparing passive drifting to DVM20 or DVM40 without nearshore retention, which have RMSD ranging from 0.44-0.91°C, and the strongest agreement for the four simulation pairs comparing DVM20 to DVM40 with or without nearshore retention, which had a RMSD ranging from 0.23-0.35°C. The strength of agreement of the four simulation pairs comparing passive drifting to DVM20 or DVM40 with nearshore retention falls between the simulations with the weakest and strongest agreement and has a RMSD ranging from 0.37-0.48°C. For Anacapa and Scripps, T_L_ has the weakest agreement for the two simulation pairs comparing passive drifting to DVM20 without nearshore retention, which had a RMSD ranging from 0.49-0.54°C and 0.72-0.80°C respectively. T_L_ shows slightly stronger agreement for the four simulation pairs comparing passive drifting to DVM40 and DVM20 to DVM40 without nearshore retention with a RMSD ranging from 0.35-0.48°C for Anacapa and from 0.43-0.53°C for Scripps. T_L_ shows the strongest agreement for the six simulations pairs that included nearshore retention with a RMSD ranging from 0.24-0.34°C for Anacapa and from 0.34-0.43°C for Scripps. ME for T_L_ did not show consistent patterns as a function of behavior, nearshore retention, or recruitment site.

#### Source site temperature

Source site temperature (T_S_) is defined as the temperature that adult urchins at the source sites are exposed to during the reproductive period of September to November. Since the reproductive period occurs in the fall while spawning and recruitment occurs in the following spring, the modeling period for T_S_ is limited to 17 years from 1996-2012. T_S_ for the baseline simulation of passive drifting, a static PLD, and no nearshore retention is shown in Fig. 3(g,h) for Gaviota and Scripps. For all months and recruitment sites, T_S_ for the baseline simulation ranges from 14.2-20.5°C. Mean baseline T_S_ over the modeling period is 17.2°C for Gaviota, 17.3°C for Ellwood, 17.4°C for Stearns, 17.2°C for Anacapa, and 17.5°C for Scripps. For all years except 1998, the baseline T_S_ decreases over the reproductive period with the change in T_S_ ranging from 0.5°C to 4°C and averaging 2.4°C. When baseline T_S_ at Gaviota is compared to the other recruitment sites, T_S_ is found to have a strong linear correlation for all sites with *r* ranging from 0.97 to 1.0. Baseline T_S_ at Gaviota is similar in magnitude to Ellwood, Stearns, and Anacapa with RMSD ranging from 0.14-0.25°C. The largest difference in magnitude of baseline T_S_ is between Gaviota and Scripps with a RMSD of 0.49°C, reflecting the mean north-south temperature gradient in the SC Bight (Fig. 5(c)).

##### Planktonic larval duration

The sensitivity testing of T_S_ to PLD shows strong agreement for simulation pairs with static and dynamic PLD for all recruitment sites. For the six simulation pairs tested for PLD, the RMSD ranges from 0.02-0.22°C, the *r* ranges from 0.99-1.00, and the ME ranges from 0.01-0.10°C.

##### Nearshore retention

Testing the sensitivity of T_S_ to nearshore retention shows strong linear correlations for all simulation pairs and recruitment sites with *r* ranging from 0.91-1.00, but displays a range of differences in magnitude as a function of behavior and site location. For the SBC mainland sites, the sensitivity testing of T_S_ to nearshore retention shows the weakest agreement for the two passive drifting simulation pairs, which produce RMSD ranging from 0.68-1.1°C, and the strongest agreement for the four DVM20 and DVM40 simulation pairs, which produce RMSD from 0.19-0.45°C. The ME for the SBC mainland sites shows that that T_S_ for the passive drifting simulations with nearshore retention is smaller by 0.53-0.91 °C than the passive drifting simulations without nearshore retention and that T_S_ for the DVM simulations with nearshore retention is smaller by 0.02-0.33 °C than the DVM without nearshore retention. For Anacapa, the sensitivity testing of T_S_ to nearshore retention shows a similar but less pronounced pattern as the SBC mainland sites with the two passive drifting simulation pairs producing RMSD from 0.43-0.45°C and the four DVM20 and DVM40 simulation pairs producing RMSD from 0.17-0.27°. The ME for Anacapa shows that that T_S_ for the passive drifting simulations with nearshore retention is smaller by 0.24-0.30 °C than the passive drifting simulations without nearshore retention and that T_S_ for the DVM simulations with nearshore retention is smaller by 0.05-0.0.8 °C than the DVM without nearshore retention. For Scripps, sensitivity testing of T_S_ to nearshore retention does not show changes in magnitude that correlate with behavior and produces RMSD from 0.38-0.52°C for all simulation pairs. The ME for Scripps shows that that T_S_ for all simulations with nearshore retention are greater by 0.31-0.40 °C than all simulations without nearshore retention, the opposite of the SBC sites.

##### Larval behavior

Testing the sensitivity of T_S_ to behavior shows strong linear correlations for all simulation pairs and recruitment sites with *r* ranging from 0.86-0.99 and differences in magnitude as a function of behavior, nearshore retention, and site location. For the SBC mainland sites, sensitivity testing of T_S_ to behavior shows the weakest agreement for the four simulation pairs comparing passive drifting to DVM20 or DVM40 without nearshore retention, producing RMSD from 0.97-1.2°C. T_S_ for the SBC mainland sites shows the strongest agreement for the four simulation pairs comparing DVM20 to DVM40 with and without nearshore retention, producing RMSD from 0.07-0.31°C. T_S_ agreement for the four simulations pairs comparing passive drifting to DVM20 or DVM40 with nearshore retention fall in between the simulations pairs with the strongest and weakest agreement, producing RMSD from 0.32-0.48°C. The ME for the SBC mainland sites shows that that T_S_ for the DVM simulations without nearshore retention is smaller by 0.73-0.95°C than the passive drifting simulations without nearshore retention and that T_S_ for the DVM simulations with nearshore retention is smaller by 0.24-0.39 °C than the passive drifting simulations with nearshore retention. The ME of 0.01-0.12°C suggests that there is little difference in T_S_ between DVM20 and the DVM40 simulations with and without nearshore retention.

For Anacapa and Scripps, agreement of T_S_ between simulations pairs is function of larval behavior and nearshore retention similar to the SBC mainland sites, but with overall stronger agreement between the simulation pairs than the SBC mainland sites. T_S_ for the four simulation pairs comparing passive drifting to DVM20 or DVM40 without nearshore retention produces the weakest agreement with RMSD ranging from 0.60-0.67°C for Anacapa and from 0.28-0.33°C for Scripps. T_S_ for the four simulation pairs comparing DVM20 to DVM40 with and without nearshore retention produces the strongest agreement with RMSD ranging from 0.13-0.32°C for Anacapa and from 0.05-0.18°C for Scripps. T_S_ for the four simulation pairs comparing DVM20 to DVM40 with nearshore retention produces medium agreement with RMSD ranging from 0.32-0.25°C for Anacapa and from 0.16-0.22°C for Scripps. For Anacapa, the ME shows that that T_S_ for the DVM simulations is smaller than T_S_ for the passive drifting simulations by 0.43-0.47°C without nearshore retention and by 0.22-0.26 C with nearshore retention. For Scripps, the ME shows that the T_S_ for the DVM simulations with and without nearshore retention is smaller than T_S_ for the passive drifting simulations with and without nearshore retention by 0.11-0.18°C. For both Anacapa and Scrips, the ME of 0.01-0.03°C suggests that there is little difference in T_S_ between the DVM20 and the DVM40 simulations with and without nearshore retention.

### 3.2 Comparison of ocean condition indices to larval recruitment index

To determine the influence of ocean conditions on larval recruitment, the 12 simulations for each ocean condition index are linearly correlated to a larval recruitment index for all recruitment sites. A meaningful linear correlation between an ocean condition index and the larval recruitment index should have strong agreement or high values of *r*. A northern SC Bight recruitment index is used for the Gaviota, Ellwood, Stearns, and Anacapa sites, which has five poor recruitment years of 1998 and 2004-2007 and 13 average or above-average recruitment years. A southern SC Bight recruitment index is used for the Scripps site, which has 13 poor recruitment years and five average and above-average recruitment years of 1996 and 1999-2002.

#### Larval dispersal distance

The strength of correlation between LDD simulations and the larval recruitment index varies as a function of larval behavior and recruitment site. For the SBC sites, LDD for all passive drifting simulations have weak and inconsistent correlations with *r* ranging from −0.23 to 0.24. For the SBC mainland sites, LDD for the DVM20 simulations show a weak but consistent inverse correlation with *r* ranging from −0.15 to −0.28. For Anacapa, LDD for the DVM20 simulations has very weak correlations with *r* ranging from −0.07 to 0.0. For the SBC sites, LDD for the DVM40 simulations have the strongest correlations with *r* ranging from −0.18 to −0.54. For Scripps, LDD has similarly strong and consistently inverse correlations for all simulations with *r* ranging from - 0.48 to −0.73. Although ranging in correlation strength, the consistent presence of negative *r* for the DVM20 and DVM40 simulations for the SBC mainland and Scripps sites suggests that LDD is inversely proportional to the larval recruitment index and that shorter LDD is correlated with higher recruitment.

#### Source zone strength

For SZS, zone combinations are identified that display a positive correlation with the larval recruitment index, implying that more larvae originate from these zones in average and above-average recruitment years than in poor recruitment years. For all sites, simulations, and zone combinations, SZS displays a very wide range of positive *r* from 0.01-0.76. The two simulations with DVM20 behavior, no nearshore retention, and a static or dynamic PLD display the strongest correlations across all sites with *r* ranging from 0.48-0.55 for the SBC sites and from 0.67-0.70 for the Scripps site. The zones associated with these correlations are Zones 1 and 3 for the SBC sites and Zones 6, 7 and 8 for the Scripps site. For the SBC sites, SZS is higher for Zone 1 than Zone 3, but the difference between Zones 1 and 3 is greater for the SBC mainland sites than Anacapa. For the SBC mainland sites, the mean SZS for the modeling period ranges from 59-70% for Zone 1 and 11-12% for Zone 3. For Anacapa, the mean SZS ranges from 35-38% for Zone 1 and 20-21% for Zone 3. For Scripps, SZS is higher for Zone 6 than Zones 7 and 8 and is lower for Zones 6-8 than Zone 1 for the SBC sites. The mean SZS for the modeling period ranges from 22-30% for Zone 6, 15-17% for Zone 7, and 12-17% for Zone 8.

To display the linear correlation between SZS and the larval recruitment index, Fig. 6(a,b) shows plots of SZS versus larval recruitment index for the simulation with DVM20, no nearshore retention, and a static PLD for Gaviota and Scripps. Using Zones 1 and 3 for Gaviota, this simulation has a *r* value of 0.55. Zone 1 is located on the SBC mainland and contains Gaviota, and Zone 3 is located on the southern coast of the North Channel Islands (Fig. 1). As shown in Fig. 6(a) for Gaviota, the SZS has a mean of 80% and a standard deviation of 23% for average and above-average recruitment years and a mean of 54% and a standard deviation of 23% in the poor recruitment years. SZS for 1998 is an outlier with poor recruitment and high SZS. Using Zones 6, 7, and 8 for Scripps, the simulation with DVM20, no nearshore retention, and a static or dynamic PLD has a *r* value of 0.70. Zone 7 contains the Scripps site and is located on the southern most part of the SC Bight mainland. Zone 6 is located directly north of Zone 7 along the SC Bight mainland. Zone 8 contains the Catalina Island coastline. As shown in Fig. 6(b) for Scripps, the SZS has a mean of 81% and a standard deviation of 20% for average and above-average recruitment years and a mean of 54% and a standard deviation of 23% for the poor recruitment years.

**Figure 6:**
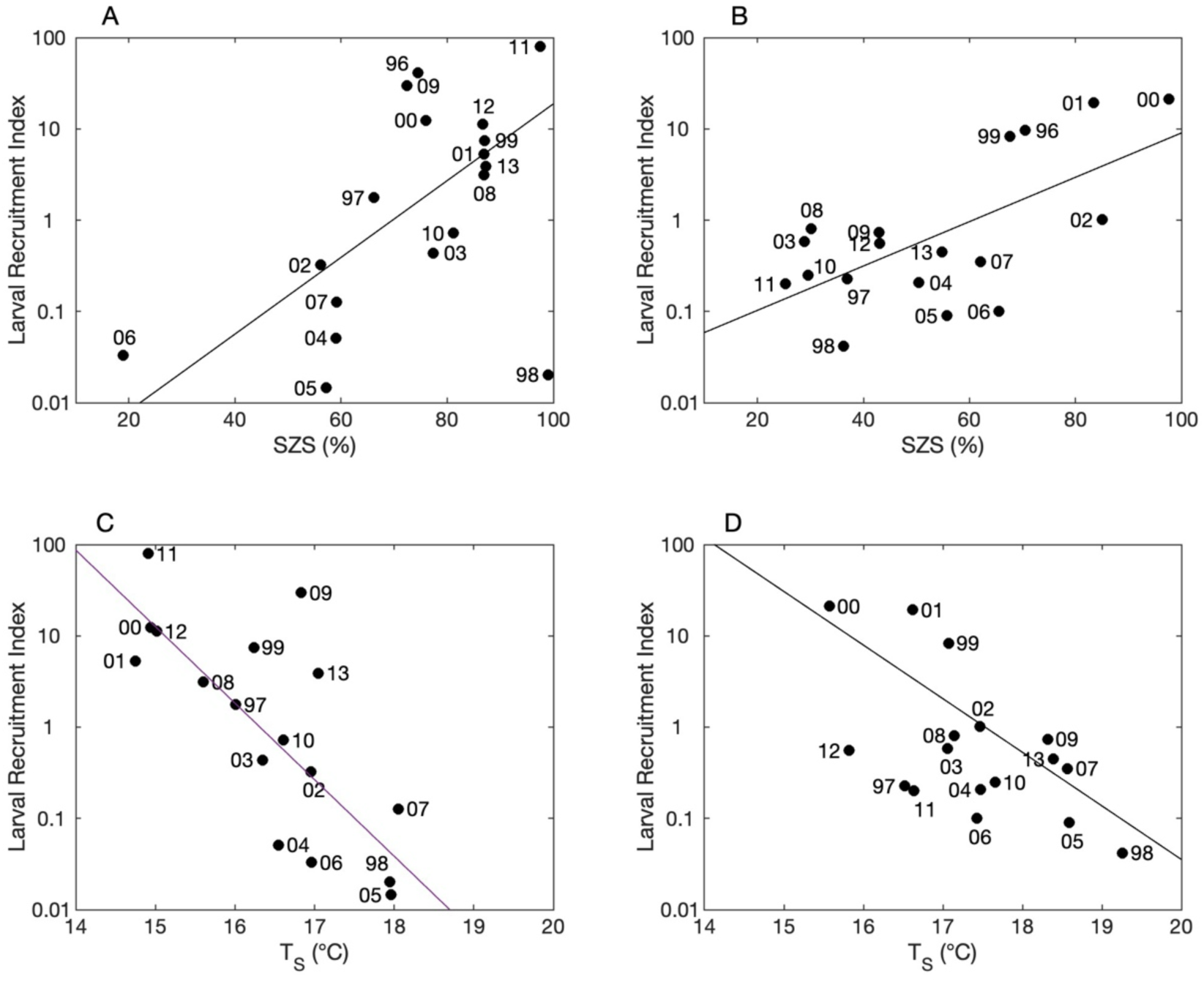
SZS (%) versus the larval recruitment index for (A) Gaviota (Zones 1 and 3) and (B) Scripps (Zones 6, 7, and 8). T_S_ (°C) versus the larval recruitment index for (C) Gaviota and (D) Scripps. The simulation shown has DVM20 behavior, no nearshore retention, and a static PLD. Each solid black circle represents one year. The solid black line shows the linear regression for all years.

#### Food supply

Comparing Chl to the larval recruitment index produces only a weak and inconsistent linear correlations for all recruitment sites and simulations. For Chl, *r* ranges from −0.52 to 0.05 with a mean of −0.13 for all simulations and recruitment sites. These results imply that that larval food supply is unlikely to be driver of larval recruitment of purple sea urchins in the SC Bight.

#### Larval temperature

When T_L_ is compared to the larval recruitment index, only weak and inconsistent linear correlations are found across all recruitment sites and simulations. For T_L_, *r* ranges from −0.35 to 0.29 with a mean of −0.13 for all simulations and recruitment sites. These results suggest that that larval temperature is not a likely driver of larval recruitment of purple sea urchins in the SC Bight.

#### Source site temperature

Out of all the OCI, T_S_ shows the highest correlation with the larval recruitment index, producing *r* with a mean of −0.69 and a standard deviation of 0.08 for all sites and simulations. The negative *r* means that high T_S_ produces to low recruitment and that low T_S_ produces average to above-average recruitment. For the SBC mainland sites, T_S_ simulations that have DVM and/or nearshore retention produce stronger correlations with *r* ranging from −0.66 to - 0.78 than the T_S_ simulations that have passive drifting without nearshore retention with *r* ranging from −0.44 to −0.55. For Anacapa and Scripps, T_S_ produces correlations with *r* ranging from −0.58 to −0.80 that do not correlate in strength with larval behavior or nearshore retention.

To display the linear correlation between T_S_ and larval recruitment index, Fig. 6(c,d) shows plots of T_S_ versus the larval recruitment index for the simulation with DVM20, no nearshore retention, and a static PLD for Gaviota and Scripps. This simulations produces a *r* of - 0.75 for Gaviota and −0.58 for Scripps. For Gaviota, the 13 years with average to above-average recruitment correlate with lower T_S_ ranging from 14.9-16.6°C and averaging 16.0°C. Three of the poor recruitment years (1998, 2005, 2007) correlate with high T_S_ of 17.8-18.0°C, but the poor recruitment years of 2004 and 2006 deviate from the linear correlation with lower T_S_ of 16.4°C and 16.5°C respectively. For Scripps, the 13 poor recruitment years have higher T_S_, ranging from 17.6-19.9°C and averaging 18.2°C. The four years with average to above-average recruitment have lower T_S_ ranging from 15.9-17.6°C and averaging 17.1°C.

## 4.0 Discussion

This study assesses the impacts of changes in circulation, temperature, and food supply on larval recruitment of purple sea urchins. Changes in circulation are quantified using LDD and SZS, which are calculated directly from particle tracks. Temperature is quantified through T_L_ and T_S_, which are calculated from the OCM’s temperature and particle tracks. Food supply is quantified using Chl, which is calculated from surface chlorophyll from satellite data and particle tracks. To accurately model larval recruitment, the larval life history of purple sea urchins must be accurately represented in the biophysical model. Although some aspects of the larval life history of purple sea urchins are known, such as the spawning and reproductive periods and potential source sites, uncertainty remains for three important aspects of larval life history, PLD, nearshore retention, and larval behavior. To address this uncertainty, sensitivity of the five OCI to these three larval life history parameters is discussed in Section 4.1. Derived from the 12 simulations with all possible combinations of larval life history parameters, each ocean condition index is then compared to a larval recruitment index to identify the potential drivers of larval recruitment, which is discussed in Section 4.2.

### 4.1 Sensitivity of ocean condition indices to larval life history parameters

#### Planktonic larval duration

All OCI shows very strong agreement between the simulation pairs with static and dynamic PLDs. This result initially appears surprising as the static PLD of 42 days is significantly different from the dynamic PLD, which ranges from 26-56 days with a mean of 37 days and standard deviation of 8 days (Fig. S2). However, Simons et al. (2013) found that particle dispersal in the SC Bight is much more sensitive to changes in particle tracking times or PLDs when PLDs are greater than ∼20 days and concluded that the changes in particle dispersal patterns decrease with increasing PLD because the particles become more evenly mixed over increasingly larger areas. To determine if this could be true for our study, *r* is calculated between the PDM with a PLD of 42 days and daily PDMs with PLDs of 1-56 days for each month, simulation, and recruitment site, producing 4,320 values of *r* for each day from 1-56 days. As PDMs are 2D particle density matrices, PDMs are vectorized for a grid cell to grid cell calculation of *r* following the methods in Simons et al. (2013). Fig. 7 displays mean *r* ± 1 standard deviation versus PLD for all *r* values and shows that there is minimal difference between the PDMs with a 42-day PLD and PDMs with PLDs greater than ∼25 days. As all OCI are derived from the particle tracks, which are used to create the PDMs, this similarity between the PDM with a PLD of 42-days and PDMs with PLDs greater than ∼25 days explains why the OCI are relatively insensitive to the static PLD of 42 days versus the dynamic PLD with a range of 26-56 days.

**Figure 7:**
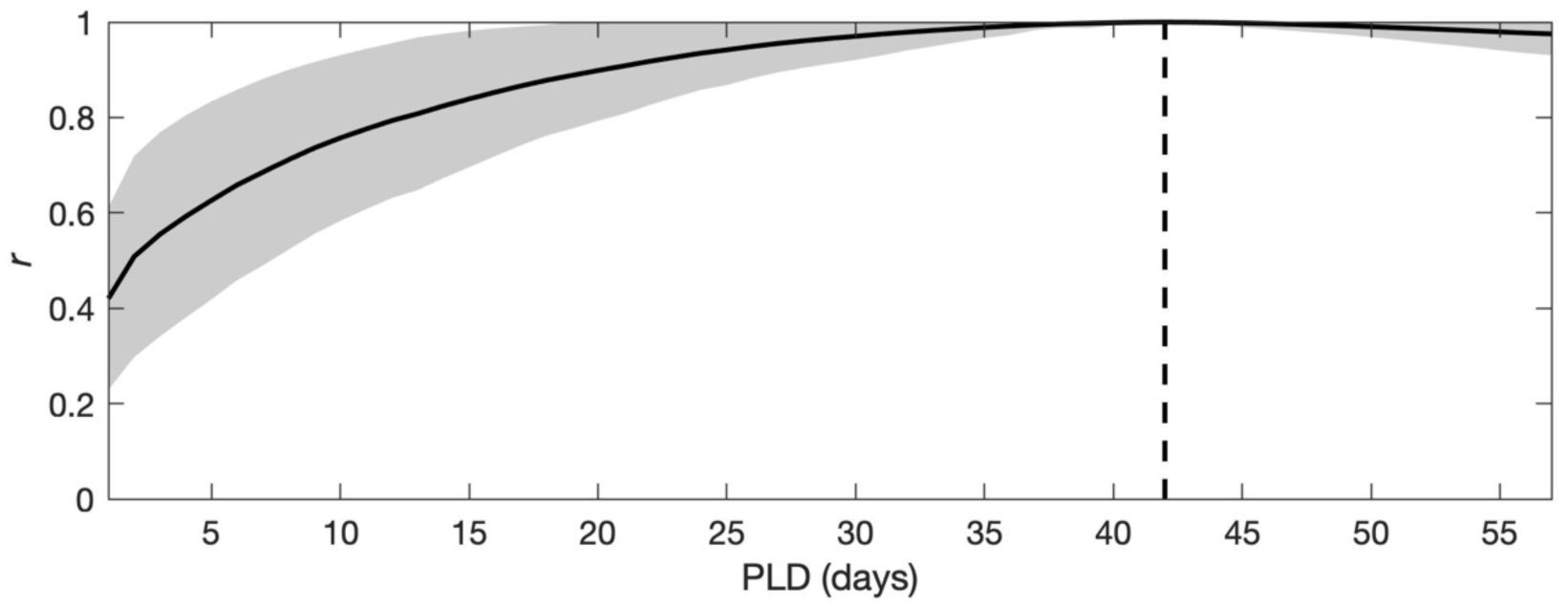
Pearson’s *r* versus PLD (days). *r* is calculated between the particle density map (PDM) with a PLD of 42 days and the PDMs with PLDs of 1-56 days for all simulations, recruitment sites, and months of the modeling period. Solid black line is mean r. Gray area is ±1 standard deviation of *r*. Dashed black line identifies a static PLD of 42 days.

#### Nearshore retention and larval behavior

Two main trends are observed in the sensitivity testing of OCI to nearshore retention and larval behavior. The first trend is that the OCI representing circulation, LDD and SZS, show much higher sensitivity to nearshore retention and larval behavior than the OCI representing food supply, Chl, and temperature, T_L_, and T_S_, for all simulations and recruitment sites. The second trend is that simulation pairs for nearshore retention and larval behavior show similar levels of sensitivity between the SBC mainland sites for all OCI and varying levels of sensitivity between the SBC mainland sites, the Anacapa site, and the Scripps sites.

Both trends appear to derive from differences in horizontal patterns of circulation, temperature, and chlorophyll within SC Bight. Fig. 5(a) shows the mean surface chlorophyll from satellite data averaged temporally over the recruitment period for the modeling period of 1996-2013. Fig. 5(b,c) displays the mean temperature from the OCM averaged vertically over the top 50 m of the water column and temporally over the recruitment and reproductive periods respectively for the modeling period. Fig. 5(d) shows the mean current from the OCM averaged vertically over the top 50 m of the water column and temporally over the recruitment period for the modeling period. The top 50 m of the water column is selected because ∼70% of the particles with passive drifting remain the top 50 m of the water column and 100% of the particles with DVM remain in the top 40 m of the water column. As shown in Fig. 5, the circulation, temperature, and chlorophyll for the areas near the recruitment sites are described below.

In the northern SC Bight, the mean circulation in the SBC, where the SBC mainland sites are located, is dominated by a cyclonic eddy, which has the strongest mean velocities in the SC Bight up to 21 cm s^-1^ and penetrates throughout the top 100 m of the water column (Oey et al. 2004; Dong et al. 2009). The strong eddy circulation creates a region of high residence times and particle accumulation in the SBC (Simons et al. 2015; Simons and Catlett 2023). The SBC is the coldest region in the SC Bight with mean temperatures 4-5°C lower than the southern SC Bight. The mean temperature immediately surrounding the SBC is ∼1.5°C higher than the mean temperature within the SBC. Driven by spring upwelling and high residence times in the SBC (Simons and Catlett 2023; Brokaw et al. 2024), the mean surface chlorophyll in the SBC is the highest in SC Bight with a maximum of 2.7 mg m^-3^ and drops to low levels of 0.1 - 1 mg m^-3^ outside of the SBC.

Located on Anacapa Island at the edge of the SBC, the Anacapa site is surrounded by the weakest mean velocities, ∼3 cm s^-1^, of the five recruitment sites and sits at the junction between flows entering the SBC and the large cyclonic eddy in the central SC Bight. The Anacapa site is located in the transition zone between the SBC and the central and southern SC Bight, where strong gradients in mean temperature and surface chlorophyll are present.

In the southern SC Bight, the circulation near the Scripps site shows southward flow along the coastline with mean velocities up to 10 cm s^-1^ and which decreases an order of magnitude to the north and east. Unlike the SBC which has consistent mean circulation to at least 100 m depth, mean circulation along the southern coastline varies with depth, displaying stronger southward flows in the top 50 m of the water column and weaker northward flows from 50-100 m depth (Dong et al. 2009). Defined by an open coastline with a narrow shelf, circulation in the region near the Scripps site produces shorter residence times and greater offshore dispersal than the SBC (Dong et al. 2012). Except for ∼5 km adjacent to the coastline, the southern SC Bight displays smaller north-south mean temperature gradients of 0.1-0.2°C per 10 km than the area surrounding the SBC of 0.3-0.4°C per 10 km. The mean surface chlorophyll throughout the southern SC Bight is very low from 0.1-1 mg m^-3^.

In order to compare the level of sensitivity between OCI, which have different units, RMSD for each ocean condition index is normalized by dividing the RMSD by the mean ocean condition index for both simulations in the pair, producing a RMSD percentage for each simulation pair. RMSD percentage is calculated for the nearshore retention and larval behavior simulation pairs with the highest sensitivity for the five OCI. For nearshore retention, the simulations pair with the highest sensitivity consists of the passive drifting simulation without nearshore retention and the passive drifting simulation with nearshore retention. For larval behavior, the simulation pair with the highest sensitivity consists of the passive drifting simulation without nearshore retention and the DVM20 simulation without nearshore retention. SZS has the highest RMSD percent for all sites and both simulation pairs, ranging from 110-210%. LDD has the next highest RMSD percent for the nearshore retention simulation pair for all sites, ranging from 30-110%, and for the behavior simulations pair for for the SBC mainland site and the Scripps site, ranging from 41-109%. RMSD percent for Chl is lower than LDD and SZS for the nearshore retention simulation pair for all sites, ranging from 21-37% and for the behavior simulations pair for the SBC mainland site and the Scripps site, ranging from 24-51%. Anacapa is the only site that has similar levels sensitivity to LDD and Chl. For all sites and both simulation pairs, T_L_ and T_S_ have the lowest RMSD percent by 1-2 orders of magnitude compared SZS, LDD, and Chl, ranging from 2-7% with a mean of 5%. For all OCI, the SBC mainland sites have similar levels of RMSD percentage, but distinctly different levels of RMSD percentage from the Anacapa and Scripps sites.

To understand why OCI and recruitment sites show different levels of sensitivity, we examine the relationship of particle dispersal to circulation, temperature, and surface chlorophyll. To quantify particle dispersal, mean PDMs for the modeling period are displayed in Fig. 2 for the Gaviota and Scripps sites for the three simulations used in the nearshore retention and larval behavior simulation pairs with the highest sensitivity. The three simulations are (1) passive drifting without nearshore retention (Fig. 2(a,d)), (2) passive drifting with nearshore retention (Fig. 2(b,e)), and (3) DVM20 without nearshore retention (Fig. 2(c,f)). All simulations shown have a static PLD. The Gaviota and Scripps sites are selected because their results reflect the difference in circulation between the SBC and the southern SC Bight.

##### Circulation (LDD and SZS) and particle dispersal

When nearshore retention is included in a simulation, the number particles coming from source sites near the recruitment site automatically increases as simulations with nearshore retention includes all particle tracks from a source site to a recruitment site that are at or below the PLD, e.g. PLDs less than or equal to 42 days. In comparison, simulations without nearshore retention only include particle tracks from a source site to a recruitment site at the specified PLD, e.g. PLDs equal to 42 days. The increase in particle concentrations near the Goleta and Scripps sites when nearshore retention is applied to the passive drifting simulation is shown in Fig. 2(a-b,d-e). As particles with shorter tracking times have a shorter dispersal distance, the LDD decreases by an average of 47% for all sites when nearshore retention is applied to the passive drifting simulation. For the same reason, the number particles in the source zones where recruitment sites are located increases when nearshore retention is applied to the passive drifting simulation. As a results, the pattern of SZS changes for all sites from being more evenly spread throughout the 9 source zones (Fig. 2(g,j)) to being dominated by the zone where the recruitment site is located (Fig. 2(h,k)). In the zones where the recruitment sites are located, the SZS increases by an average of 44% for all sites.

When DVM is included in a simulation, particle dispersal changes as new particle tracks are produced. When compared to the passive drifting simulation without nearshore retention, the DVM20 simulation without nearshore retention decreases particle dispersal for the SBC mainland sites, which results in a decrease of LDD by an average of 82%. The decrease in particle dispersal for the Gaviota site can be seen in Fig. 2(a,c). For the SBC mainland sites, long residence times in the SBC appear to limit particle dispersal for particle tracks with DVM, but not for particle tracks with passive drifting. This decrease in particle dispersal for DVM20 behavior increases particle concentrations near the SBC mainland sites and SZS by an average of 46% in Zone 1, where the recruitment sites are located. The Anacapa site shows a smaller increase in particle dispersal and LDD than the SBC mainland sites for the behavior simulation pair. Compared to the regions near the other recruitment sites, the circulation in the region of the Anacapa Island, where the Anacapa site is located, has the lowest velocities and shallowest bathymetry, which may be restricting particle dispersal regardless of larval behavior. In contrast, DVM20 behavior increases particle dispersal for the Scripps site compared to passive drifting, which results in an increase in LDD by an average of 20%. The increase in particle dispersal for DVM20 behavior appears to be driven by the open coastline and short retention times of the southern SC Bight. For the Scripps site, the increase in particle dispersal for DVM20 versus passive drifting is shown in Fig. 2(d,f). For the Scripps site, the increase in particle dispersal from DVM20 decreases the SZS for the Zone 7, where the Scripps site is located, by an average of 16% and increases the SZS for Zones 6 and 8, which are near Zone 7, by an average of 5% and 6% respectively.

##### Temperature (T_L_ and T_S_) and particle dispersal

The OCI representing temperature, T_L_ and T_S_, show the lowest sensitivity to nearshore retention and larval behavior. T_L_ is calculated from particle tracks and OCM temperature, and T_S_ is calculated from SZS and OCM temperature. Although both nearshore retention and larval behavior produce significant changes in particle dispersal, temperature is relatively consistent over the horizontal scale of the changes in particle dispersal. This can be seen by comparing the areas of high particle concentrations in the mean PDMs (Fig. 2(a-f)) with the mean temperature for the recruitment and reproductive periods in Fig. 5(b,c)). As a result, RMSD for T_L_ and T_S_ for all recruitment sites is relatively small, ranging from 0.49-0.99°C and 0.31-1.1°C respectively for the simulation pairs with the highest sensitivity. However, it is important to note that although the RMSD for T_L_ and T_S_ are small relative to their absolute values, a difference of ∼1°C could have be important if the temperature is near a threshold that is detrimental for larval development or survival.

##### Food Supply (Chl) and particle dispersal

The ocean condition index representing food supply, Chl, shows higher sensitivity than T_L_ and T_S_ and lower sensitivity than SZS and LDD for all recruitment sites. Chl is calculated directly from particle tracks and surface chlorophyll from satellite data. The highest sensitivity for Chl comes from the SBC mainland sites for the two simulation pairs comparing passive drifting with and without nearshore retention and passive drifting to DVM20 without nearshore retention. As the highest Chl in the SC Bight is confined to the SBC (Fig. 5(a)), Chl for SBC mainland sites will increase when nearshore retention increases particle concentrations in the SBC and when DVM limits particle dispersal to the SBC region (Fig. 2(c)). In contrast, Chl is much less sensitive to changes in particle dispersal outside of the SBC, including the southern SC Bight, where Chl is consistently low.

##### Summary

The higher sensitivity of LDD and SZS to nearshore retention and larval behavior than T_L_, T_S_, and Chl (trend 1) appears to be driven by the fact that circulation and subsequently particle dispersal in the SC Bight are much more horizontally variable than temperature and surface chlorophyll. As LDD and SZS are derived directly from particle tracks driven by circulation, they are more sensitive to the spatial changes in circulation than T_L_, T_S_, and Chl, which are derived from particle tracks with temperature or surface chlorophyll.

All OCI show similar levels of sensitivity between the SBC mainland sites and varying levels of sensitivity between the SBC mainland sites, the Anacapa site, and the Scripps sites (trend 2). This result appears to be driven by the strong consistent circulation in the SBC. When nearshore retention or DVM is included in a simulation for the SBC mainland sites, the circulation creates a region of high particle concentrations in the SBC, which produces similar changes in OCI for all SBC mainland sites. Due to very different circulation and retention near the Anacapa and Scripps sites, the addition of nearshore retention or DVM to the simulation for these sites does not produce changes in OCI consistent with the other recruitment sites.

The high sensitivity of LDD and SZS to nearshore retention and variable sensitivity of LDD and SZS to larval behavior as a function location within the SC Bight have important implications for the use of larval connectivity calculated from biophysical models for scientific research as well as policy decisions for environmental protection (Cowen and Sponaugle 2009; Metaxas and Saunders 2009; Swearer et al. 2019). These results show that selection of larval life history parameters can dramatically change the location of predicted larval source sites by 100s of kilometers. Thus, larval connectivity predictions that are based on a single set of larval life history parameters could provide misleading information for policy decisions, such as those for marine protected areas. For this reason, our study shows the importance of conducting sensitivity studies for larval life history parameters that can’t be derived from existing empirical data.

### 4.2 Identifying ocean conditions indices that drive of larval recruitment

Circulation, food supply, and temperature are discussed as potential drivers of larval recruitment of purple sea urchins in the SC Bight. OCI are considered potential drivers of interannual changes in larval recruitment if a strong linear correlation is found between the ocean condition index and the larval recruitment index for one or more simulations and all recruitment sites.

#### Circulation (LDD and SZS)

The OCI representing circulation, LDD and SZS, displayed a range of correlation strength as function of larval behavior. The simulations with the strongest correlations for both LDD and SZS included DVM. For simulations with DVM, LDD is found to be weakly to moderately inversely correlated to the larval recruitment index for all recruitment sites, implying that larval dispersal occurs over smaller areas in the average and above-average recruitment years than in the poor recruitment years. For the zones containing and close to the recruitment sites, SZS for the DVM20 simulations without nearshore retention shows the strongest linear correlation with the larval recruitment index, suggesting that a higher percentage of larvae come from these zones in the average and above-average recruitment years than in the poor recruitment years. From these results, we hypothesize that circulation in the SC Bight is similar among the average and above-average recruitment years and among the poor recruitment years, but different between the average and above-average recruitment years and poor recruitment years. However, our results suggest that larval recruitment is only connected to circulation when larvae have behavior, such as DVM, that limits their dispersal to the region near the recruitment sites. Although this study only compares passive drifting to two simple DVM patterns, urchin larvae in the SC Bight may performing a different behavior that also reduces larval dispersal in average and above-average recruitment years.

#### Food Supply (Chl)

Chl is used as a proxy for larval food supply in the SC Bight because it represents amount of phytoplankton present, which is the food source for sea urchin larvae. The correlation between Chl and the recruitment index is very low and inconsistent in both magnitude and sign across all simulations, implying that Chl is not driving the interannual changes in larval recruitment. For all sites, the lowest values of Chl occur during 2000, a very high recruitment year, and 1998, a very low recruitment years (Fig. 3, Table S1), implying that larvae are not food limited in the SC Bight even in years with lower primary production. It is important to note that of the five OCI in this study, Chl has the most uncertainty as it is based on surface chlorophyll and does not account for vertical changes chlorophyll.

#### Temperature (T_L_ and T_S_)

In prior work, Okamoto et al. (2020) analyzed 27-year larval recruitment datasets of purple sea urchins in the northern and southern SC Bight and found that recruitment was negatively correlated with sea surface temperature at the recruitment sites and positively correlated with the multivariate ENSO index (MEI), which tracks El Niño and La Niña events in the Pacific Ocean. Throughout the SC Bight, high temperatures and El Niño events were observed to correspond with low larval recruitment. Okamoto et al. (2020) found that recruitment was highly synchronous within the northern and southern SC Bight regions, implying that recruitment was being driven by regional trends in oceanography and not by local conditions at the recruitment sites.

The OCI that represent temperature are T_L_ and T_S_. Although Okamoto et al. (2020) found that sea surface temperature was a linearly correlation with larval recruitment, T_L_ and T_S_ show extremely different correlation strengths. T_L_ is weak and inconsistently correlated with the larval recruitment index for all sites and simulations, while T_S_ is strongly correlated with the larval recruitment index for all sites and simulations. These results imply that T_S_ is likely driving the interannual changes in larval recruitment, but T_L_ is not. To understand why T_L_ and T_S_ produce divergent results, T_L_ and T_S_ are compared to documented thresholds for larval survival and reproduction for the purple sea urchin in the SC Bight.

In the SC Bight, the threshold temperature for lethal conditions of purple sea urchin larvae increases as temperature approaches 18-20°C (Munstermann et al. In Review). T_L_ for all simulations and recruitment sites is below this threshold of ∼18°C. For the modeling period of 1996-2013, the highest recorded temperatures in the SC Bight occurred during a El Niño event that lasted from July 1997 to June 1998 (Lynn and Bograd 2002; McClatchie et al. 2016). However, during the 1998 recruitment period of March-June, T_L_ is far below the lethal level with maximum T_L_ of 15.9°C for all simulations and recruitment sites. Fig. 8 shows the mean seasonal temperature cycle at the Gaviota and Scripps recruitment sites using temperatures from the OCM averaged over the 18-year modeling period. These recruitment sites represent temperature extremes in SC Bight with Gaviota in the cooler northern region and Scripps in the warmer southern region (Fig. 5(b,c)). In Fig. 8, the recruitment period of March-June shows mean temperatures at Gaviota ranging from 12.0-14.3°C and Scripps ranging from 14.4-16.6°C. The dashed lines from March-June in Fig. 8 represent the temperatures at Gaviota and Scripps during the 1998 El Niño event over the recruitment period, which range from 12.9-16.6°C and show that even during a strong El Niño event, the temperatures during the recruitment period do not vary significantly from the mean annual cycle. Thus, strong El Niño events, although known for producing high temperatures, appear to have minimal effect on temperatures in the SC Bight during the recruitment period, remaining well below the lethal limit of 18°C. As warming produced by marine heatwaves, such as the 1997-1998 El Niño event, does not raise T_L_ above the lethal limit for urchin larvae during the recruitment period, we hypothesize that T_L_ is most likely not driver of larval recruitment for purple sea urchins in the SC Bight.

**Figure 8:**
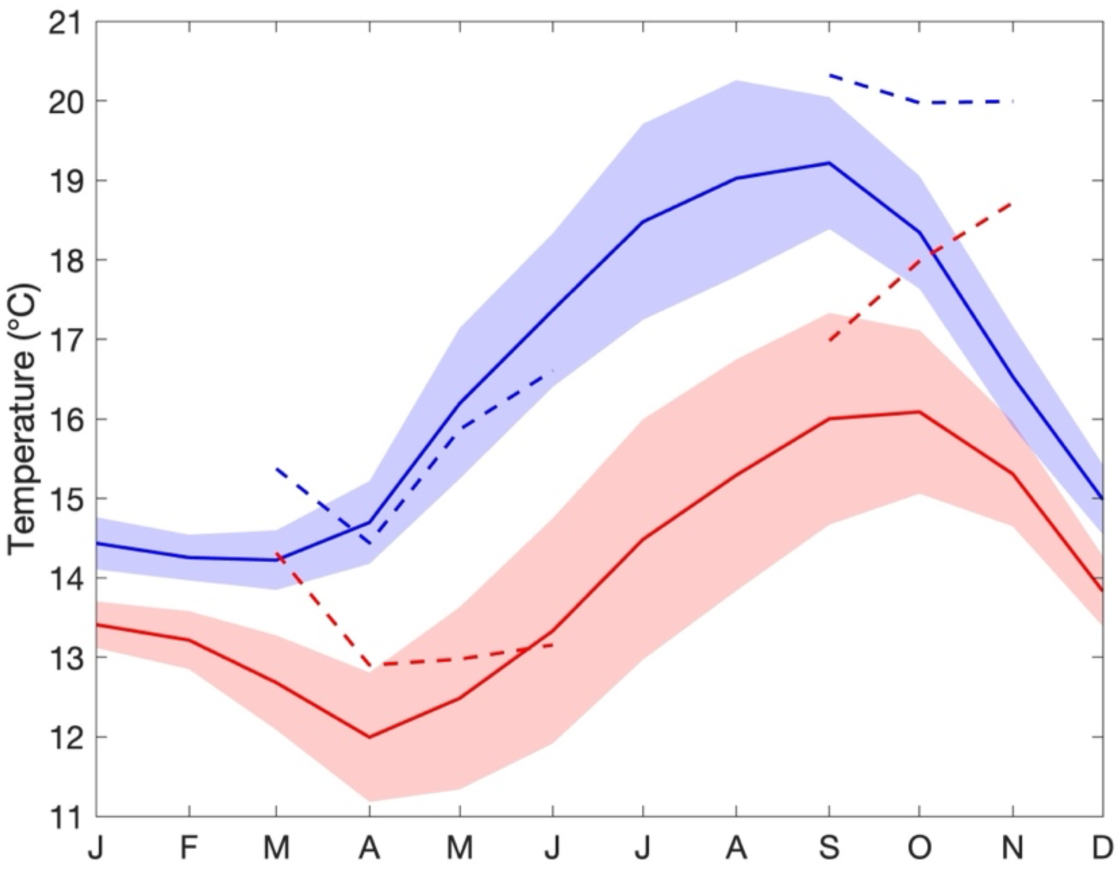
Mean annual temperature cycle ± 1 standard deviation (°C) at 5m depth for Gaviota (red) and Scripps (blue). Dashed lines represent the reproductive period (September-November) for 1997 and the recruitment period (March-June) for 1998. Temperature is derived from the ocean circulation model and is averaged over the modeling period of 1996-2013.

Okamoto et al. (Accepted) evaluated the impact of temperature on the reproduction of purple sea urchins and found that simulating historical heatwave conditions of temperatures greater than ∼18°C suppressed female reproduction. For the simulations with DVM and/or nearshore retention, T_S_ for SBC mainland sites is below the threshold of 18°C for average and above-average recruitment years with a mean and standard deviation per simulation ranging from 15.5-16.4°C and 1.1-1.3°C. For the same simulations, T_S_ for SBC mainland sites is warmer and close to the 18°C threshold for the poor recruitment years with a mean and standard deviation per simulation ranging from 17.2-17.8°C and 1.2-1.4°C. For Anacapa and Scripps, the difference in T_S_ between the average and above-average recruitment years and poor recruitment years is smaller than the SBC sites, but still shows that the good recruitment years are colder below threshold of 18°C and the poor recruitment years are warmer near or above the threshold. For Anacapa, T_S_ has a mean and standard deviation per simulation ranging from 16.3-16.6°C and 1.3-1.4°C for average and above-average recruitment years and from 17.5-17.8°C and 1.1-1.3°C for poor recruitment years. For Scripps, T_S_ has a mean and standard deviation per simulation ranging from 16.6-17.2°C and 1.0-1.3°C for average and above-average recruitment years and from 17.7-18.3°C and 1.5-1.6°C for poor recruitment years and. These results imply that T_S_ is a likely driver of larval recruitment and that recruitment along the SBC mainland is more strongly tied to T_S_ than recruitment near Anacapa Island and the southern SC Bight mainland.

The SBC mainland sites display the strongest correlation between T_S_ and the larval recruitment index for the simulations with DVM and/or nearshore retention. DVM and nearshore retention both increase the concentrations of particles in Zone 1, where the SBC mainland sites are located, which consequently increases the SZS for Zone 1 and more heavily weights T_S_ by the temperature close to the recruitment sites. This result implies that for the SBC mainland sites, larval sources in near the recruitment sites are more likely driving recruitment at these sites than larval sources in zones farther away from the recruitment sites. For the Anacapa and Scripps sites, the inclusion of DVM and nearshore retention in the simulations did not produce a higher correlation between T_S_ and the larval recruitment index, which suggest that the larval sources in the zone nearest these sites may not be as important in driving larval recruitment than the SBC mainland sites.

Okamoto et al. (2020) also found that the larval recruitment index correlated strongly with MEI. To explore this correlation, *r* is calculated for mean T_S_ and mean MEI over the reproductive period. Yearly mean T_S_ and MEI are found have a strong positive linear correlation with *r* ranging from 0.63-0.82 with a mean of 0.81 for all simulations and sites. These results suggest that El Niño events, represented by a positive MEI, are correlated with high T_S_ and poor recruitment. The strongest El Niño event of the modeling period occurred from July 1997 to June 1998 and had a mean MEI of 2.1 over this time period. In Fig. 8, the dashed lines from September to November represent the temperatures at the Gaviota and Scripps sites during the 1997 reproductive period and show that the temperatures in the SC Bight were significantly higher than the annual average. Fig. 9(a) shows the larval recruitment index as a function of T_S_ versus MEI for the simulation with DVM20, no nearshore retention, and a static PLD for all recruitment sites. The data in Fig. 9(a) suggests that T_S_ greater than ∼17.5°C and MEI greater than ∼-0.5 are likely to produce poor larval recruitment. Fig. 9(b) shows the larval recruitment index as a function of T_S_ versus SZS for the Zones 1 and 3 for the SBC sites and Zones 6, 7, and 8 for the Scripps site for the simulation with DVM20, no nearshore retention, and a static PLD. Fig. 9(b) suggests that in average recruitment year, T_S_ is less than ∼17°C and the SZS for the zones near the recruitment site is greater than ∼60%. In Fig. 9(b), 1998 for the SBC mainland sites is an outlier as it is a poor recruitment year, but shows high SZS for the Zones 1 and 3. SZS for the 1998 recruitment period correlates with T_S_ for 1997 reproductive period. T_S_ in 1997 is the highest of the 18-yr modeling period, ranging from 17.9-18.2°C, for the SBC mainland sites and the only year where monthly T_S_ does not decrease over the reproductive period (Fig. 3(g)). Thus for 1998, we hypothesize that although the circulation in the SBC correlates with average to above-average recruitment, the high temperatures during the 1997 reproductive period reduced larval supply in 1998 to the extent that circulation is no longer a factor in recruitment.

**Figure 9:**
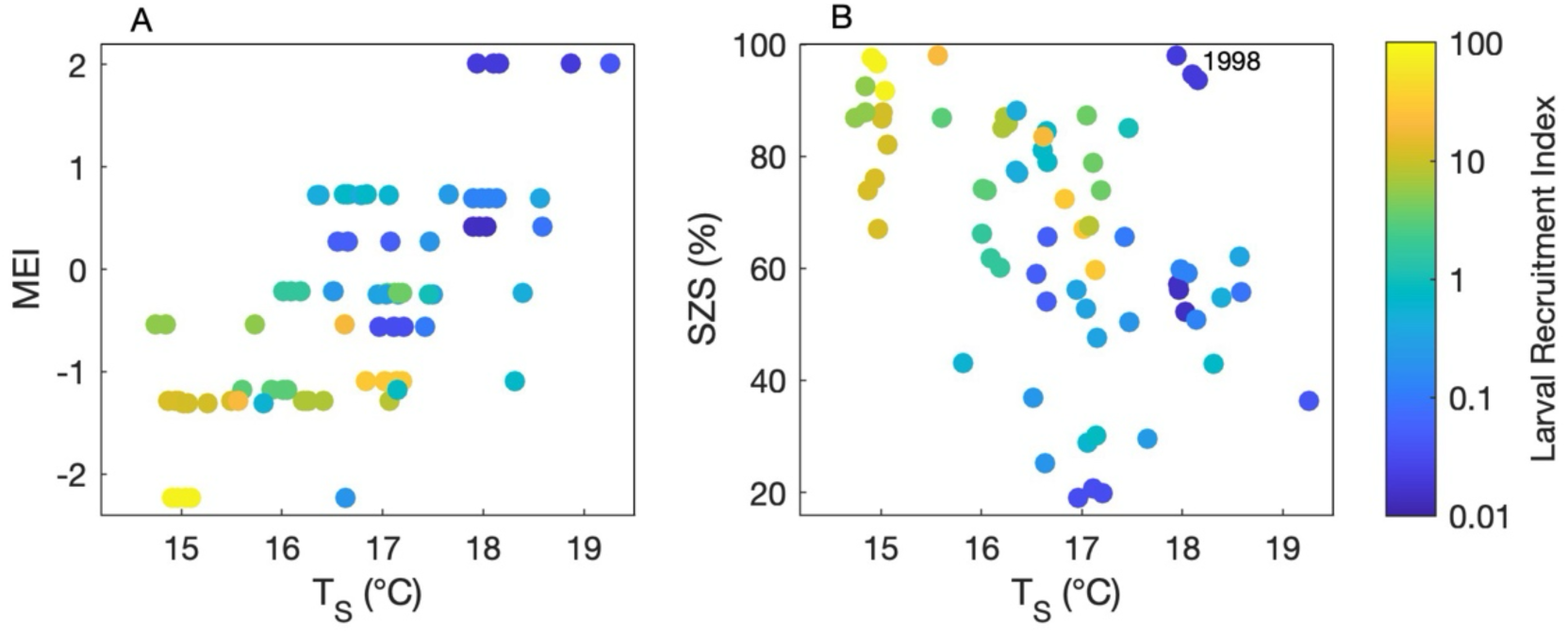
Larval recruitment index as a function of (a) T_S_ (°C) and MEI for all recruitment sites and (b) T_S_ (°C) and SZS (%) for Gaviota, Ellwood, Stearns, and Scripps. All data show is for the simulation with DVM20 behavior, no nearshore retention, and a static PLD. A circle represents one year. SZS is calculated from Zones 1 and 3 for Gaviota, Ellwood, and Stearns and Zones 6, 7, and 8 for Scripps.

#### Summary

T_S_ is hypothesized to be a primary driver of the interannual variation in larval recruitment of purple sea urchins in the SC Bight. Our results suggest that mean T_S_ over the three-month reproductive period greater than 17-18°C produces poor recruitment by suppressing reproduction in adult sea urchins at the source sites, which is supported by experimental data (Okamoto et al. Accepted). Circulation, through LDD and SZS, is found to be a potential driver of larval recruitment if the larvae have behavior that allows them to remain 35-45 km from a recruitment site inside the SBC and 80-100 km from a recruitment site in the central and southern SC Bight. As T_S_ for the SBC mainland sites had a stronger correlation with larval recruitment for the simulations that included larval life history parameters that limited dispersal distance, we hypothesize that urchin larvae in the SBC are likely behaving in a way which reduces their dispersal distance and consequently that urchin larvae are more likely to come from sources near the recruitment sites. Many prior studies attribute the settlement success to larvae coming from areas near their spawning sites due to migration and swimming behavior, even in strong upwelling systems (Tapia and Pineda 2007; Morgan 2014; Morgan et al. 2018; James et al. 2023; Meyer et al. 2024).

We found that T_S_ during the reproductive period of September to November is likely an important driver of larval recruitment, but T_L_ during the recruitment period of March to June is not, even in years with marine heat waves and strong El Niño events. These results suggest that the timing of species recruitment and reproduction is a critical factor in predicting the impact of temperature on larval recruitment. We hypothesize that marine heatwaves which occur during time of year when temperatures are at their annual maximum are more likely to produce temperatures high enough to negatively impact recruitment and reproduction than those which occur during the time of year when temperatures are at their annual minimum. Species that recruit or reproduce during the time of year when the system temperatures are at their annual maximum are more susceptible to marine heat waves than those that recruit or reproduce at other times of the year. As the frequency of marine heatwaves increases (Sydeman et al. 2014; Di Lorenzo and Mantua 2016; Bograd et al. 2023), understanding the relationship between the timing of marine heatwaves and species recruitment and reproduction becomes even more important.

## Acknowledgements

This research was supported by the National Science Foundation Biological Oceanography Program (OCE-2023693 and OCE-2023649).

