## Supplemental Information for "Impacts of changing ocean circulation, temperature, and food supply on larval recruitment of purple sea urchins in Southern California: A biophysical modeling study"

| YEAR | NORTH | SOUTH |
|---|---|---|
| 1996 | 1.62 | 0.98 |
| 1997 | 0.25 | -0.64 |
| 1998 | -1.69 | -1.38 |
| 1999 | 0.87 | 0.92 |
| 2000 | 1.09 | 1.32 |
| 2001 | 0.73 | 1.28 |
| 2002 | -0.49 | 0.00 |
| 2003 | -0.36 | -0.23 |
| 2004 | -1.29 | -0.68 |
| 2005 | -1.84 | -1.05 |
| 2006 | -1.48 | -1.00 |
| 2007 | -0.90 | -0.46 |
| 2008 | 0.49 | -0.09 |
| 2009 | 1.47 | -0.13 |
| 2010 | -0.14 | -0.60 |
| 2011 | 1.90 | -0.70 |
| 2012 | 1.05 | -0.26 |
| 2013 | 0.59 | -0.35 |

**Table S1:** Larval recruitment index for Northern (North) and Southern (South) Southern California Bight from standardized annual scale trends (Okamoto et al. 2020).

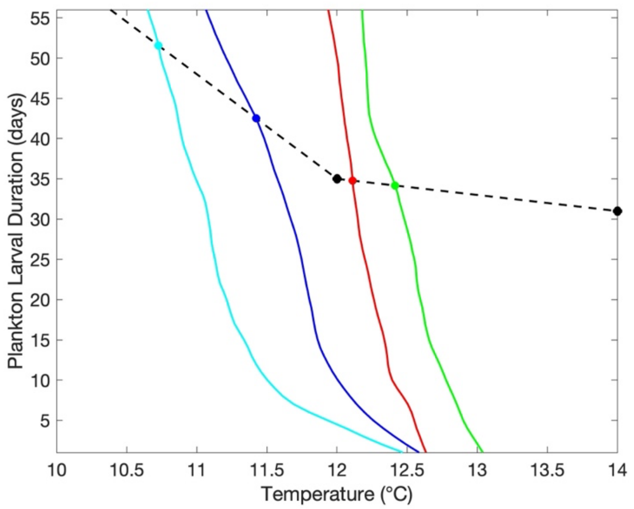


**Figure S1:** Example derivation of dynamic planktonic larval duration (PLD): PLD (days) versus mean particle temperature (°C) for the Stearns site. Red line is March 2003. Green line is April 2003. Dark blue line is May 2003. Light blue line is June 2003. The solid red, green, dark blue, and light blue circles show the dynamic PLDs of 35 days, 34 days, 43 days, and 52 days for March, April, May, and June respectively. The black dashed line and solid circles show the experimental data (Munstermann et al. 2024).

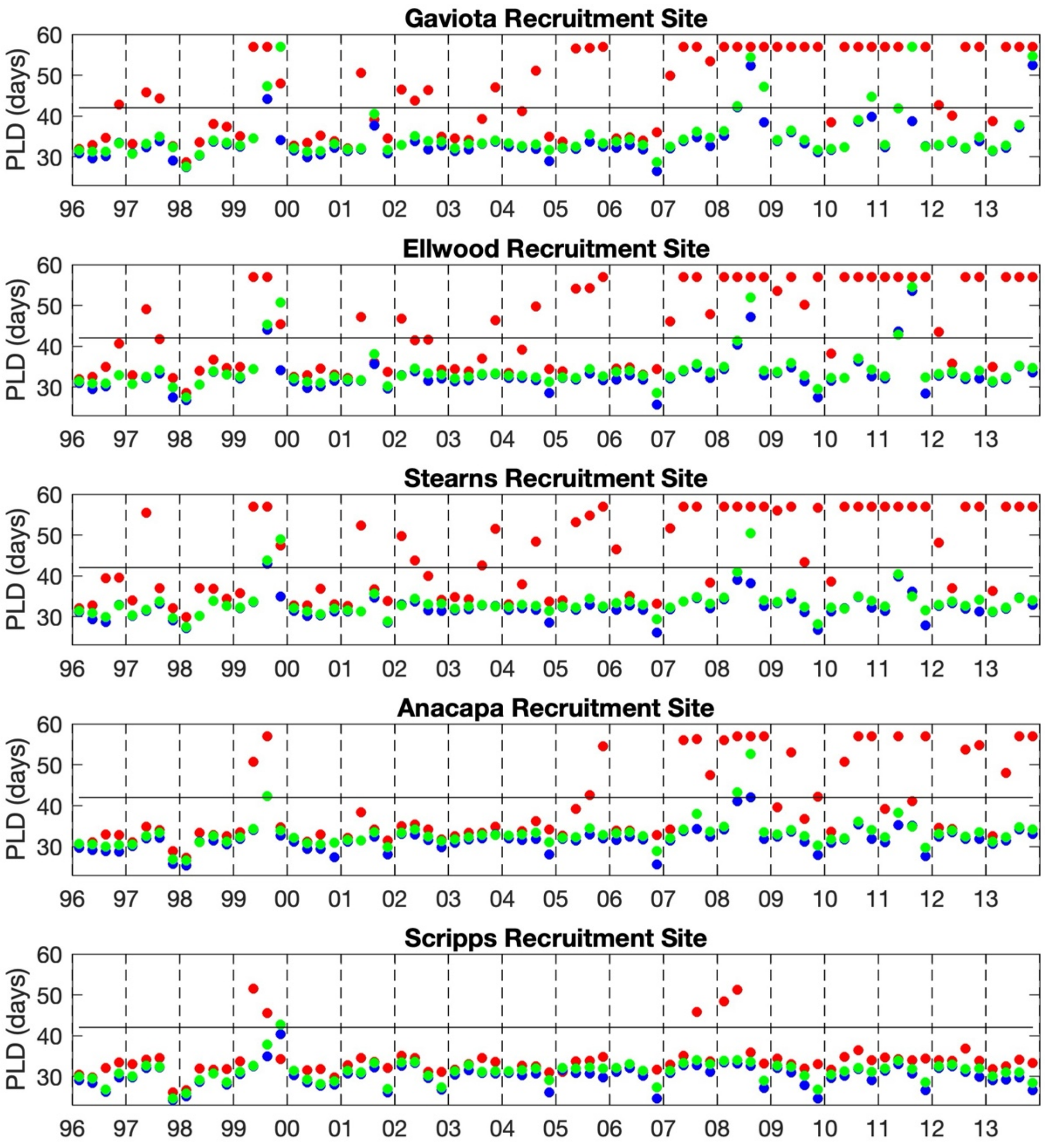


**Figure S2:** Dynamic planktonic larval duration (PLD, days) shown monthly for recruitment period of March-June. Solid red circles show passive drifting behavior. Solid blue circles show DVM20 behavior. Solid green circles shown DVM40 behavior.

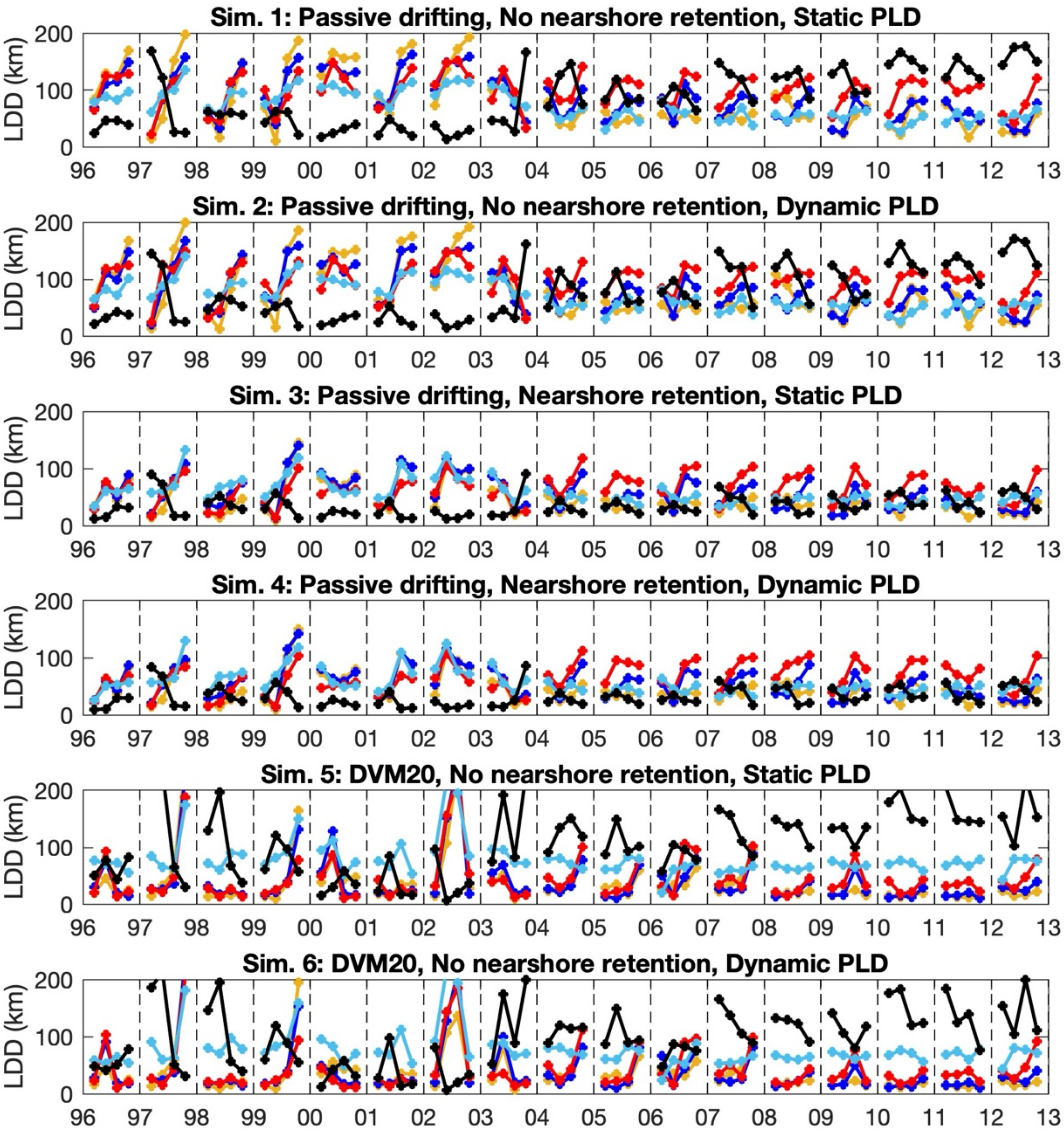


**Figure S3:** Larval dispersal distance (LDD, km) for Simulations 1-6. LDD is shown for the larval recruitment period of March-June. Gaviota is the yellow line. Ellwood is the dark blue line. Stearns is the red line. Anacapa is the light blue line. Scripps is the black line.

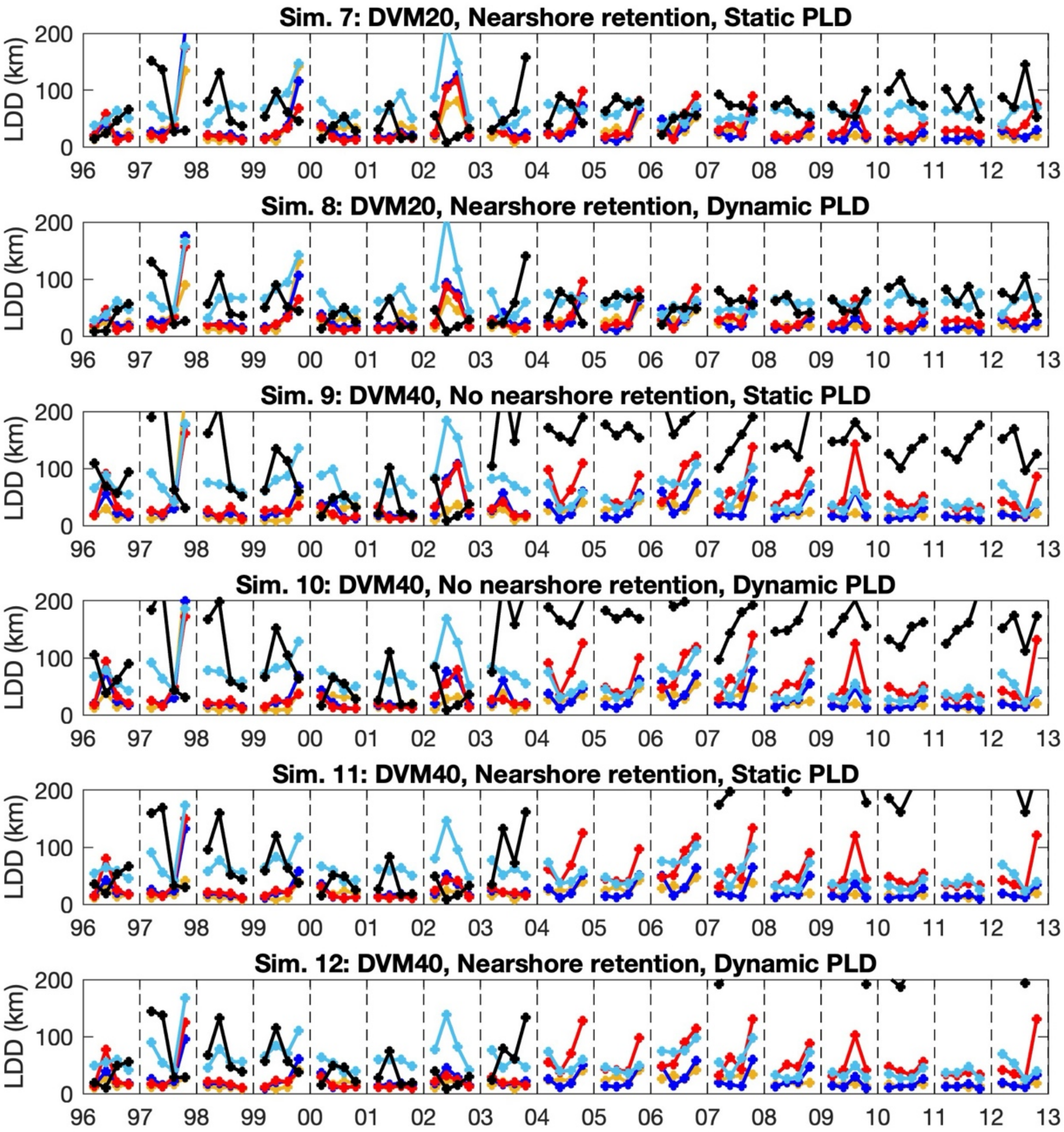


**Figure S3 (con't):** Larval dispersal distance (LDD, km) for Simulations 7-12. LDD is shown for the larval recruitment period of March-June. Gaviota is the yellow line. Ellwood is the dark blue line. Stearns is the red line. Anacapa is the light blue line. Scripps is the black line.

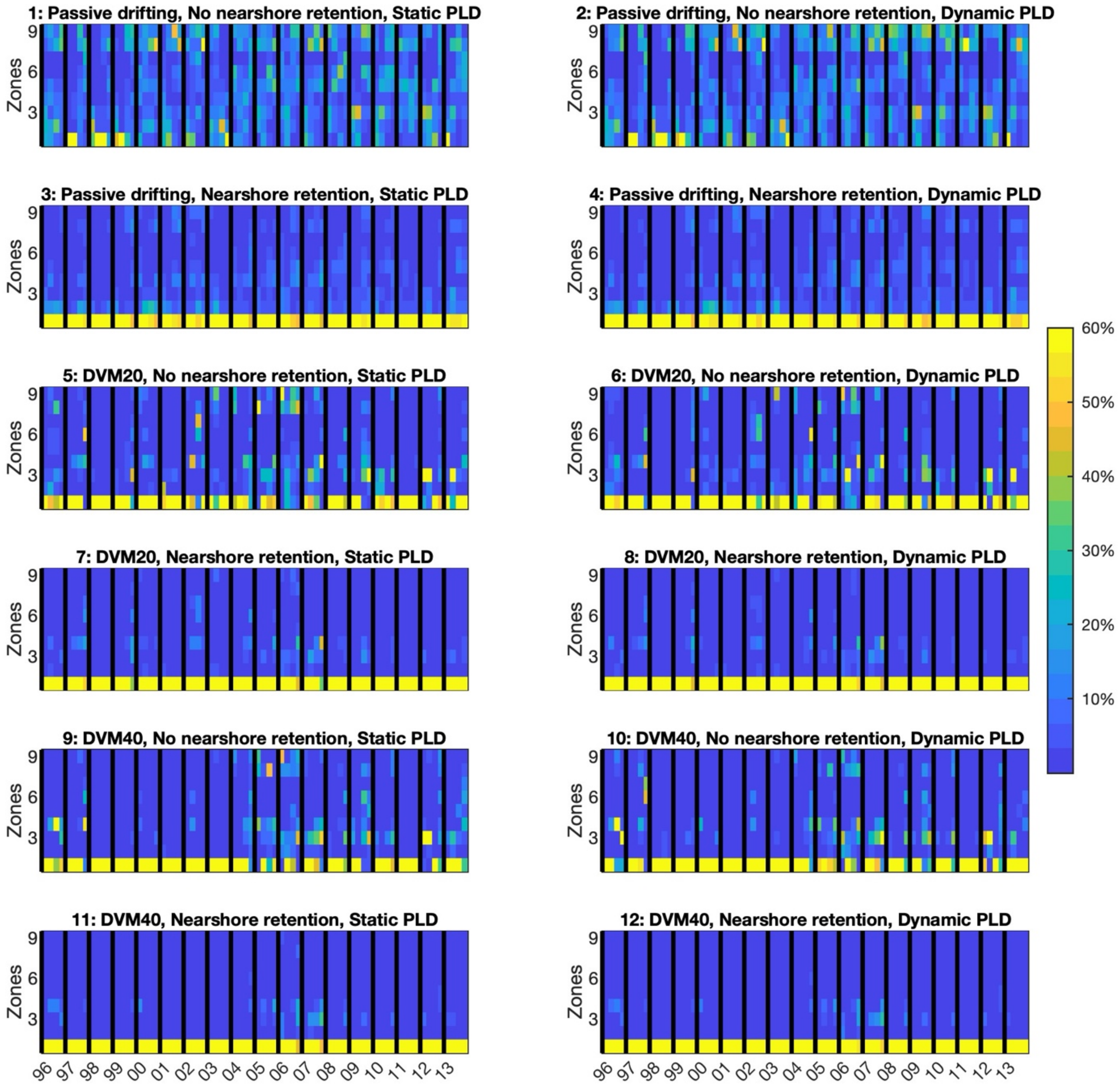


**Figure S4:** Source zone strength (SZS,%) for the Gaviota Site for Simulations 1-12. SZS is shown for the larval recruitment period of March-June. Units for SZS are in percentage of particles.

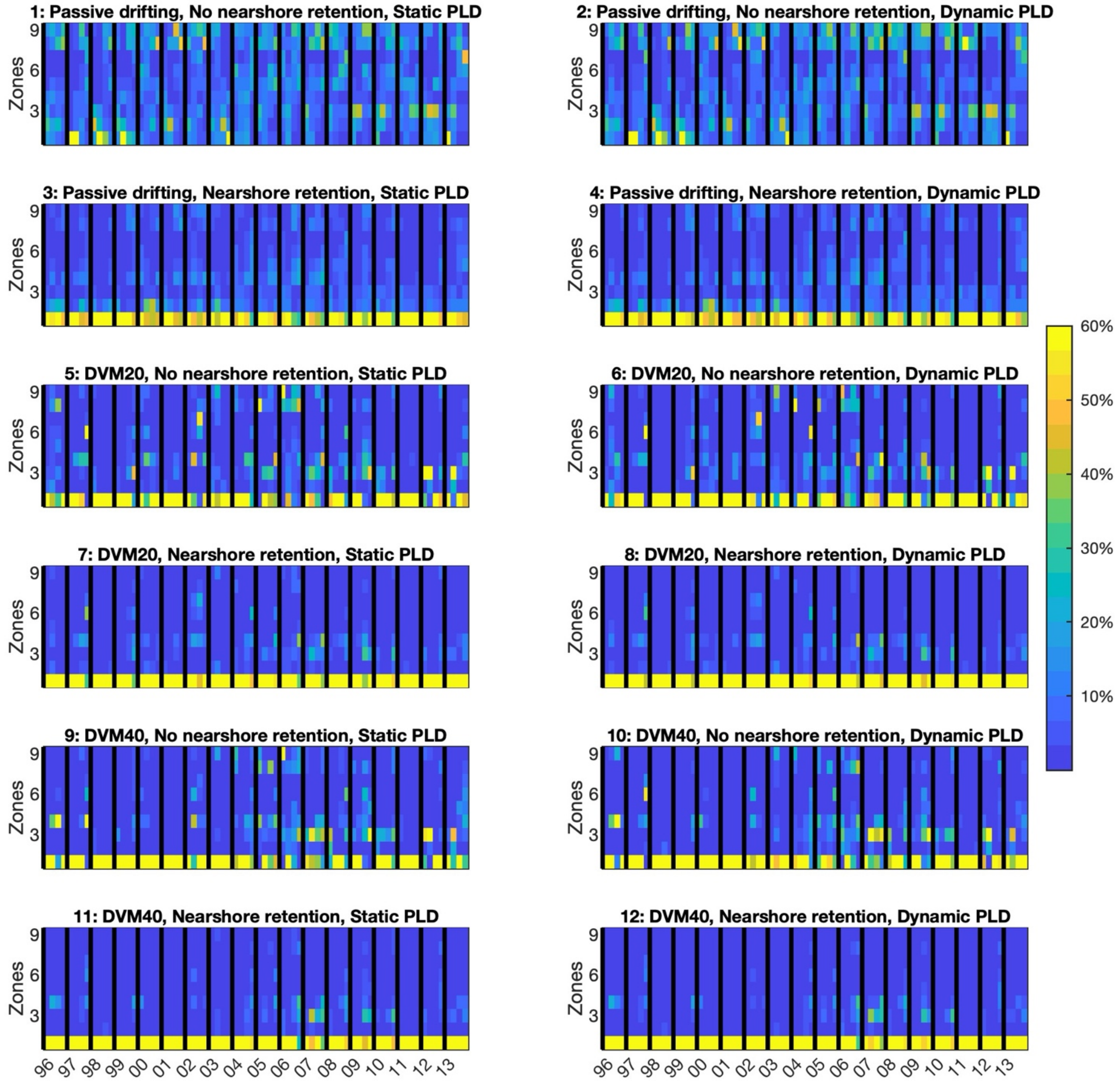


**Figure S5:** Source zone strength (SZS,%) for the Ellwood Site for Simulations 1-12. SZS is shown for the larval recruitment period of March-June. Units for SZS are in percentage of particles.

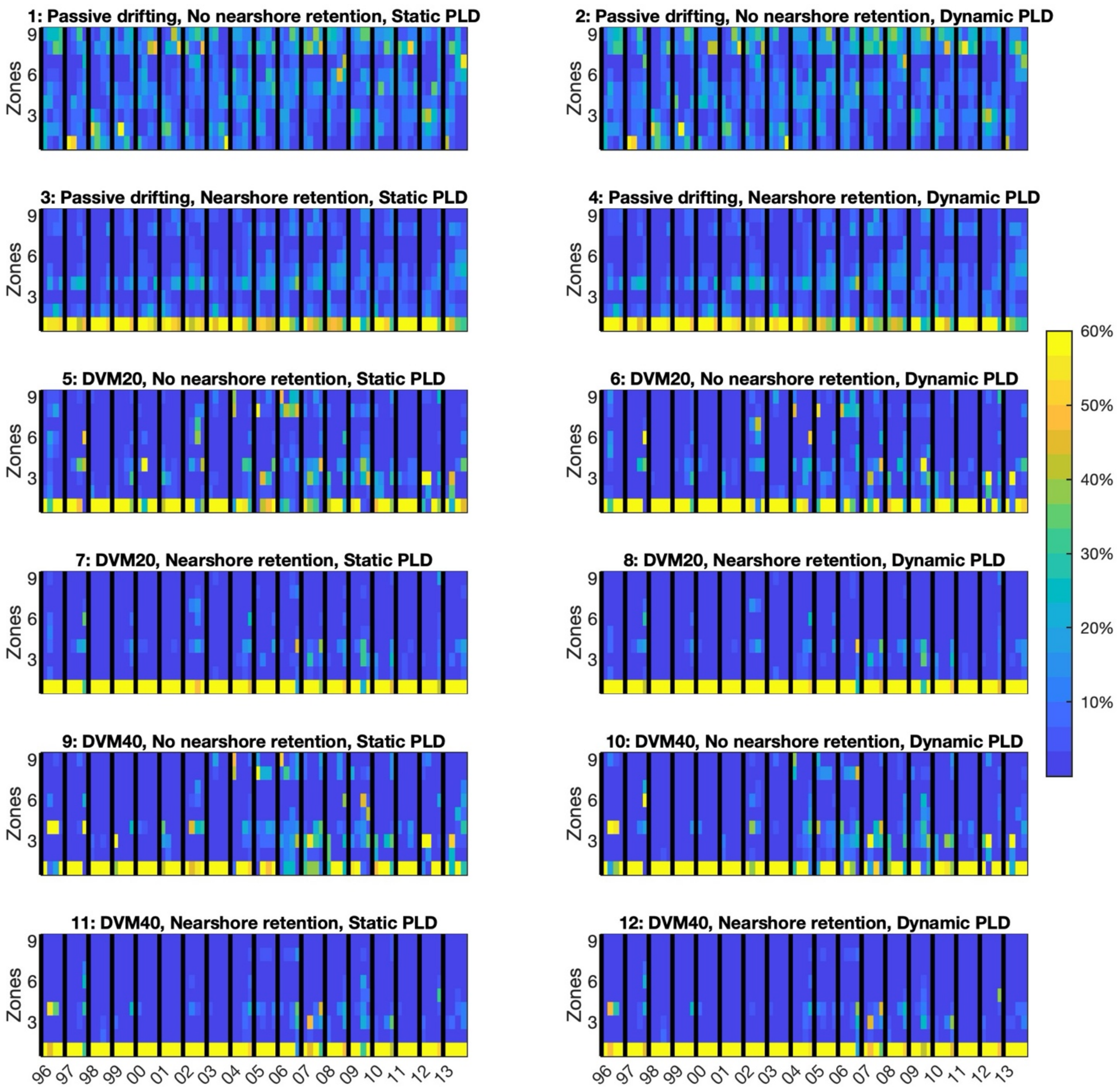


**Figure S6:** Source zone strength (SZS,%) for the Stearns Site for Simulations 1-12. SZS is shown for the larval recruitment period of March-June. Units for SZS are in percentage of particles.

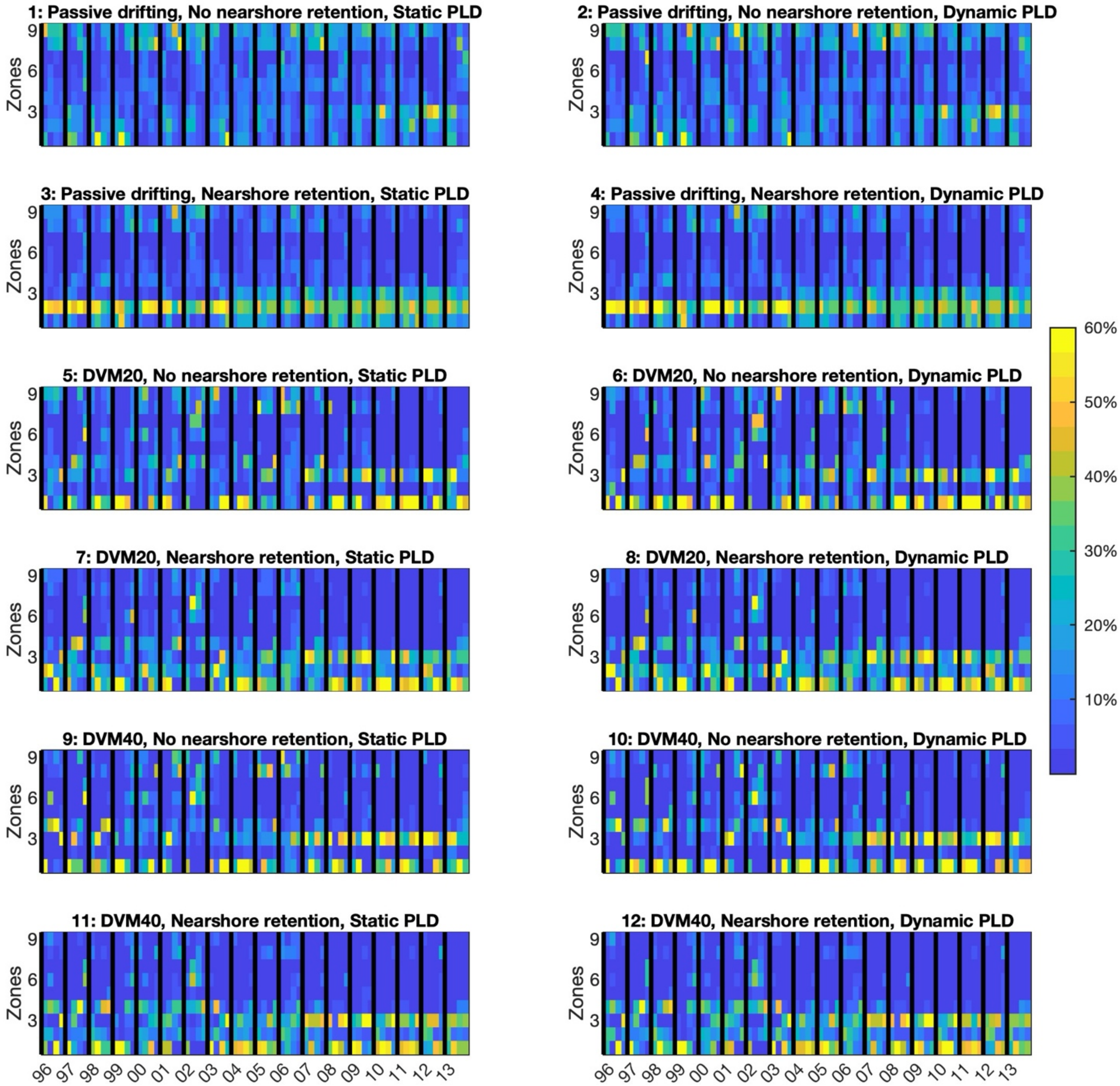


**Figure S7:** Source zone strength (SZS,%) for the Anacapa Site for Simulations 1-12. SZS is shown for the larval recruitment period of March-June. Units for SZS are in percentage of particles.

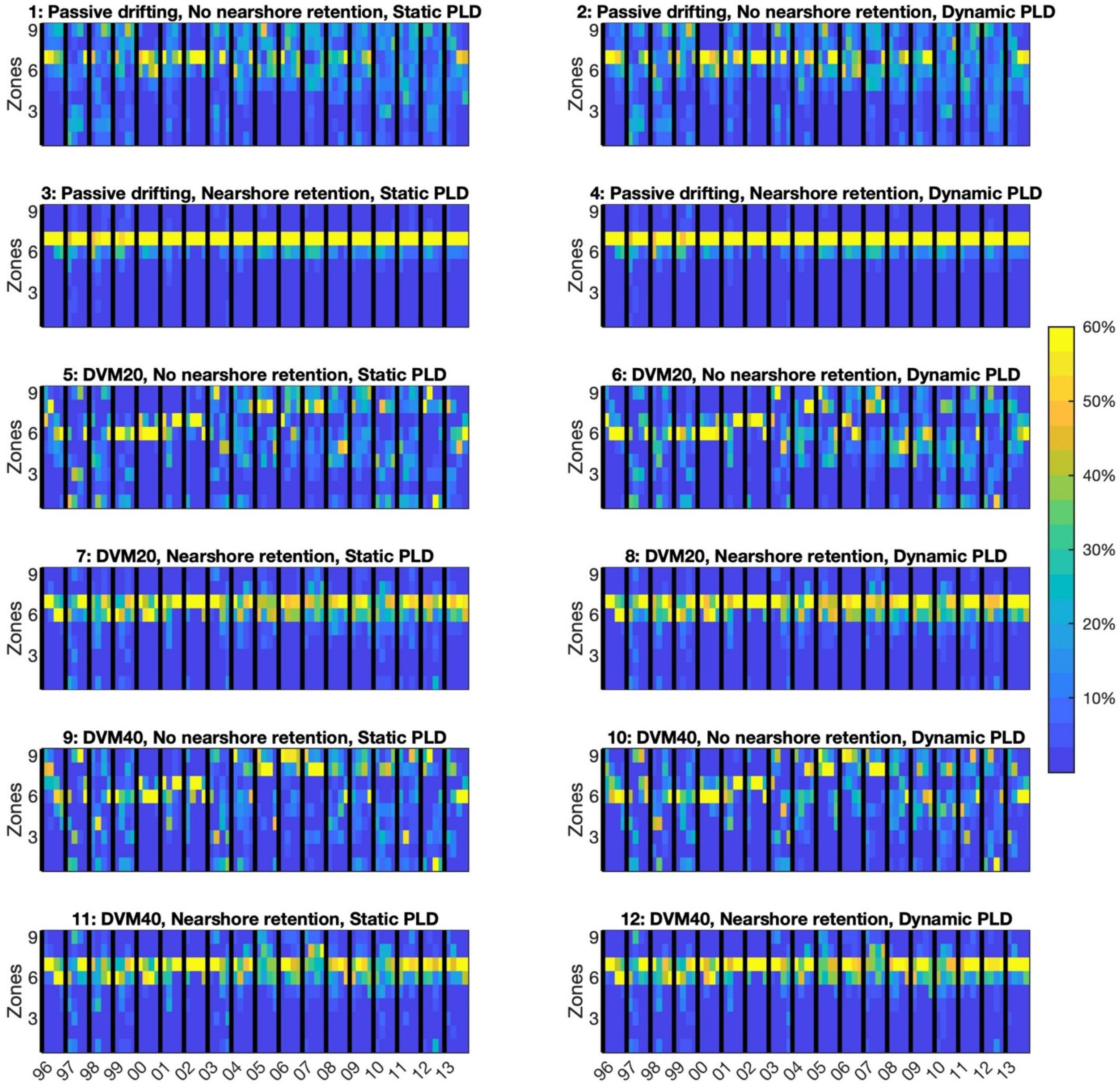


**Figure S8:** Source zone strength (SZS,%) for the Scripps Site for Simulations 1-12. SZS is shown for the larval recruitment period of March-June. Units for SZS are in percentage of particles.

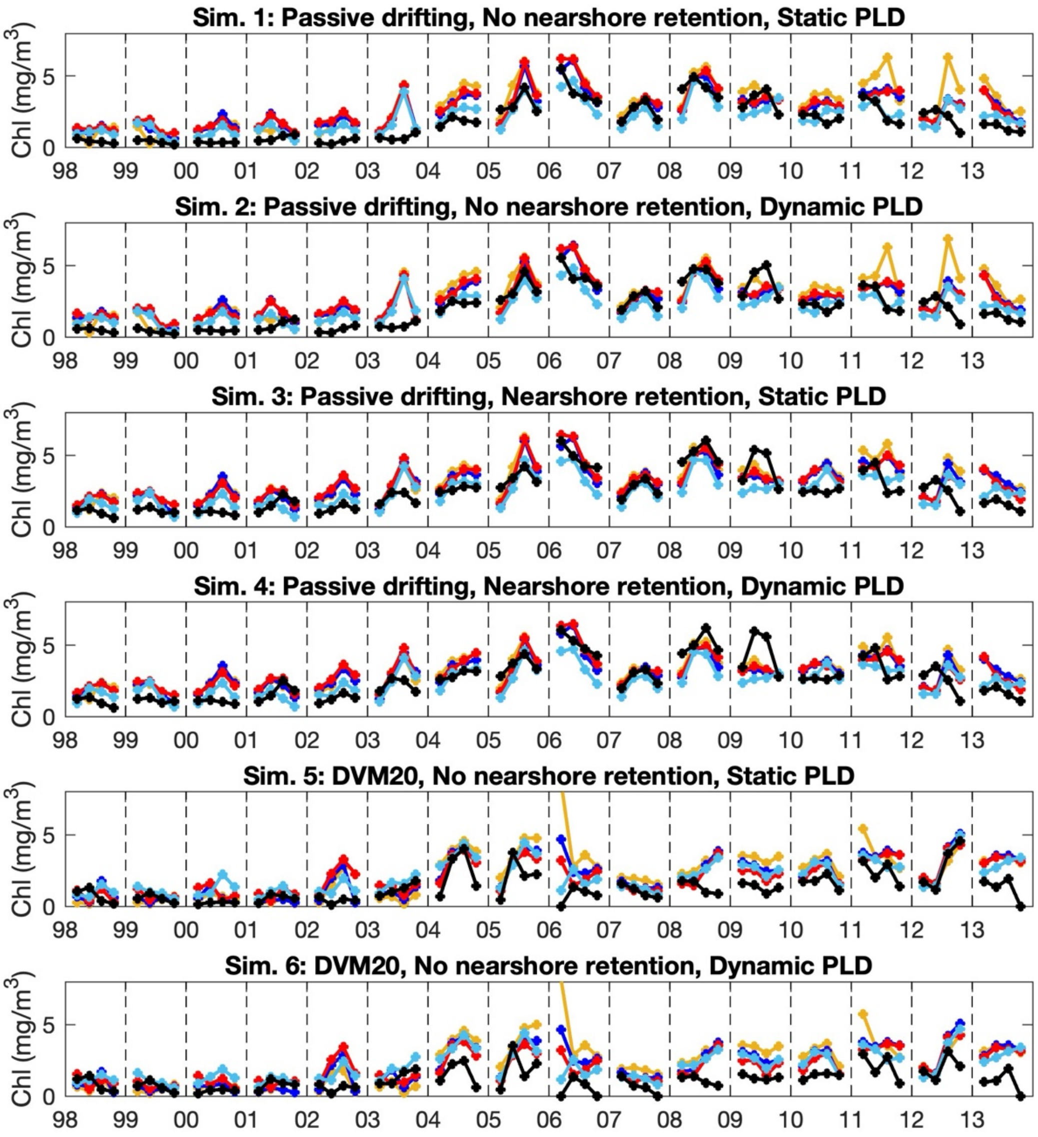


**Figure S9:** Food supply (Chl, mg $m^{-3}$) for Simulations 1-6. Chl is shown for the larval recruitment period of March-June. Gaviota is the yellow line. Ellwood is the dark blue line. Stearns is the red line. Anacapa is the light blue line. Scripps is the black line.

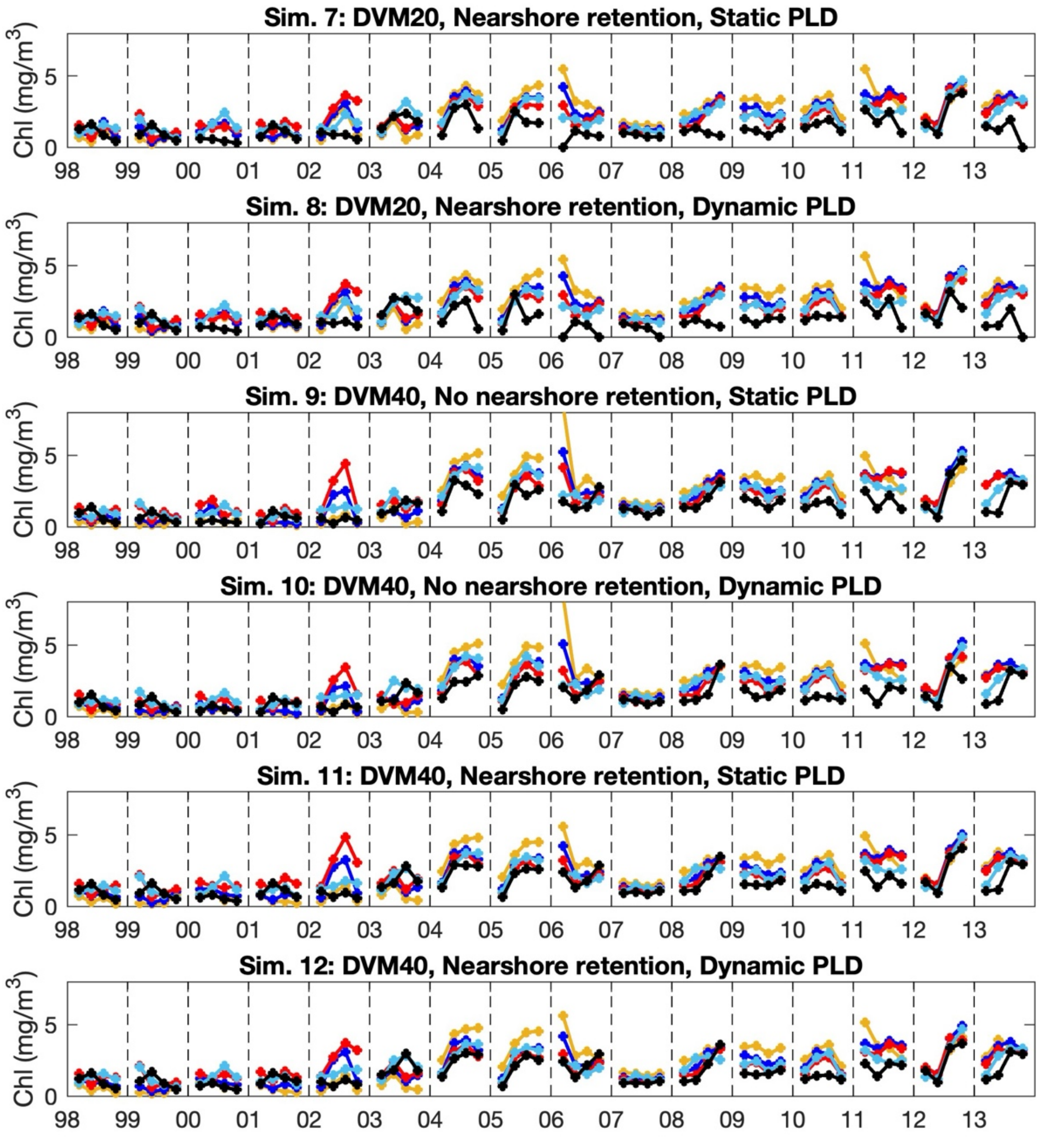


**Figure S9 (con't):** Food supply (Chl, mg m$^{-3}$) for Simulations 7-12. Chl is shown for the larval recruitment period of March-June. Gaviota is the yellow line. Ellwood is the dark blue line. Stearns is the red line. Anacapa is the light blue line. Scripps is the black line.

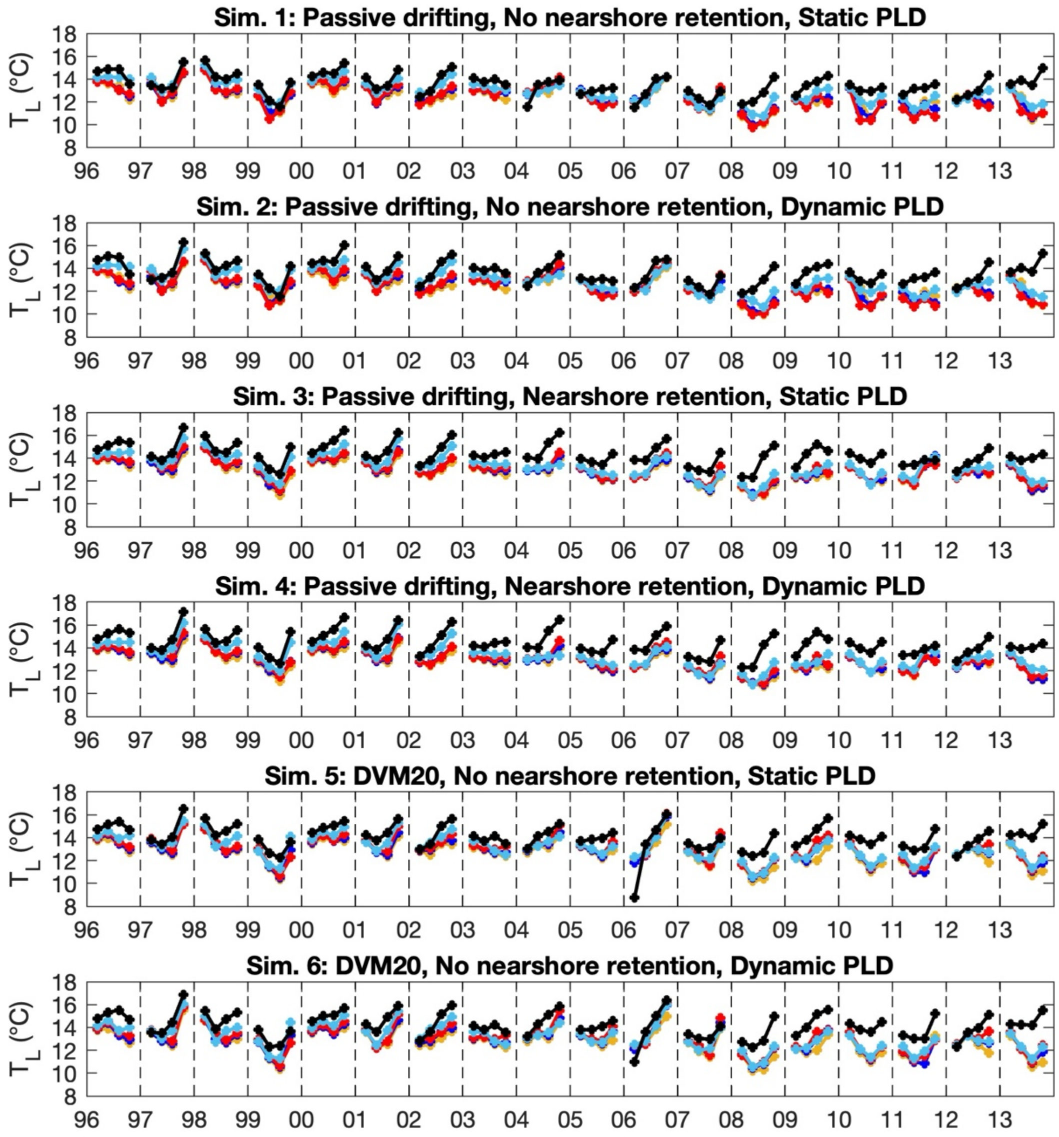


**Figure S10:** Larval temperature ($T_L$,°C) for Simulations 1-6. $T_L$ is shown for the larval recruitment period of March-June. Gaviota is the yellow line. Ellwood is the dark blue line. Stearns is the red line. Anacapa is the light blue line. Scripps is the black line.

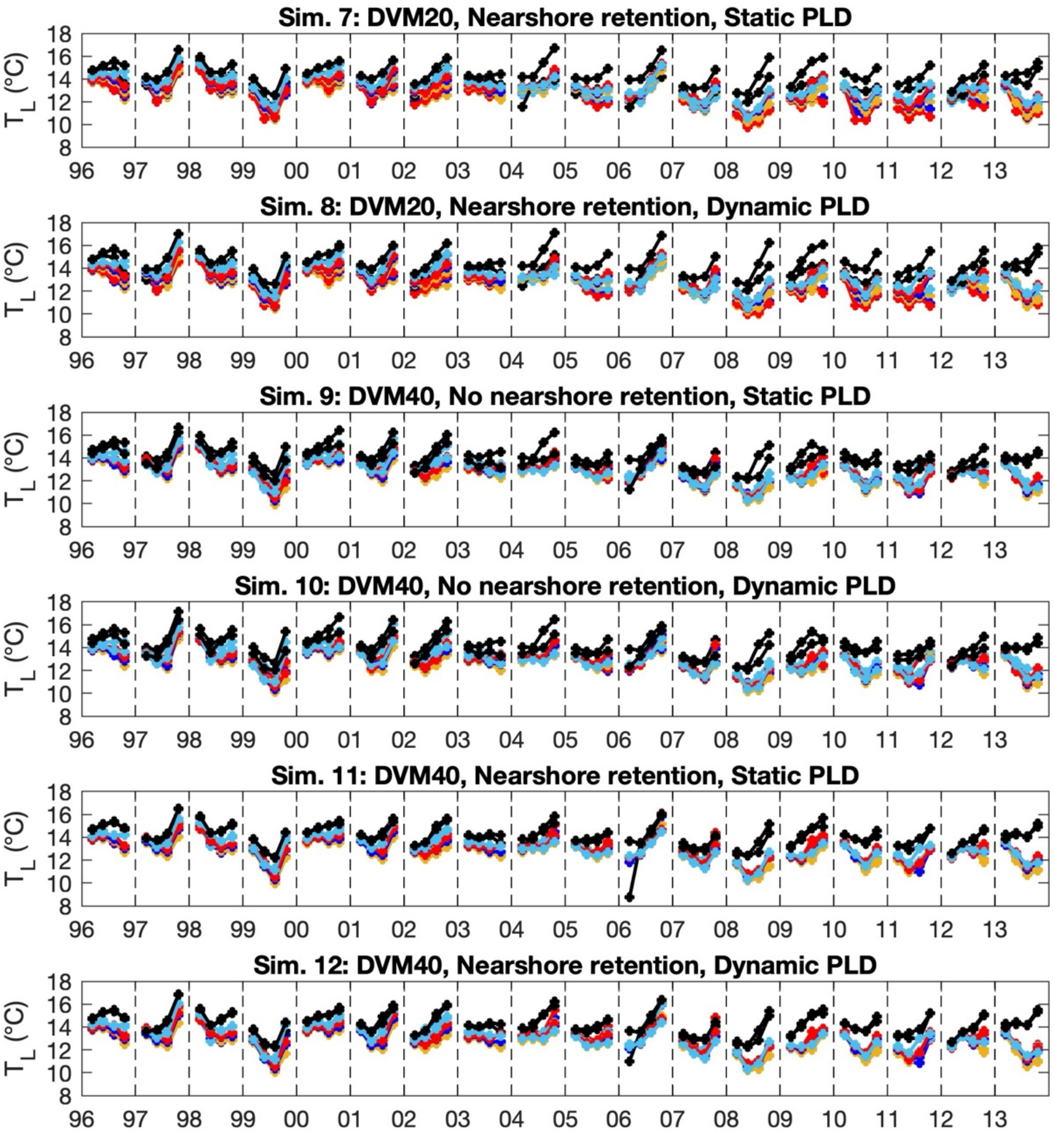


**Figure S10(con't):** Larval temperature ($T_L$,°C) for Simulations 7-12. $T_L$ is shown for the larval recruitment period of March-June. Gaviota is the yellow line. Ellwood is the dark blue line. Stearns is the red line. Anacapa is the light blue line. Scripps is the black line.

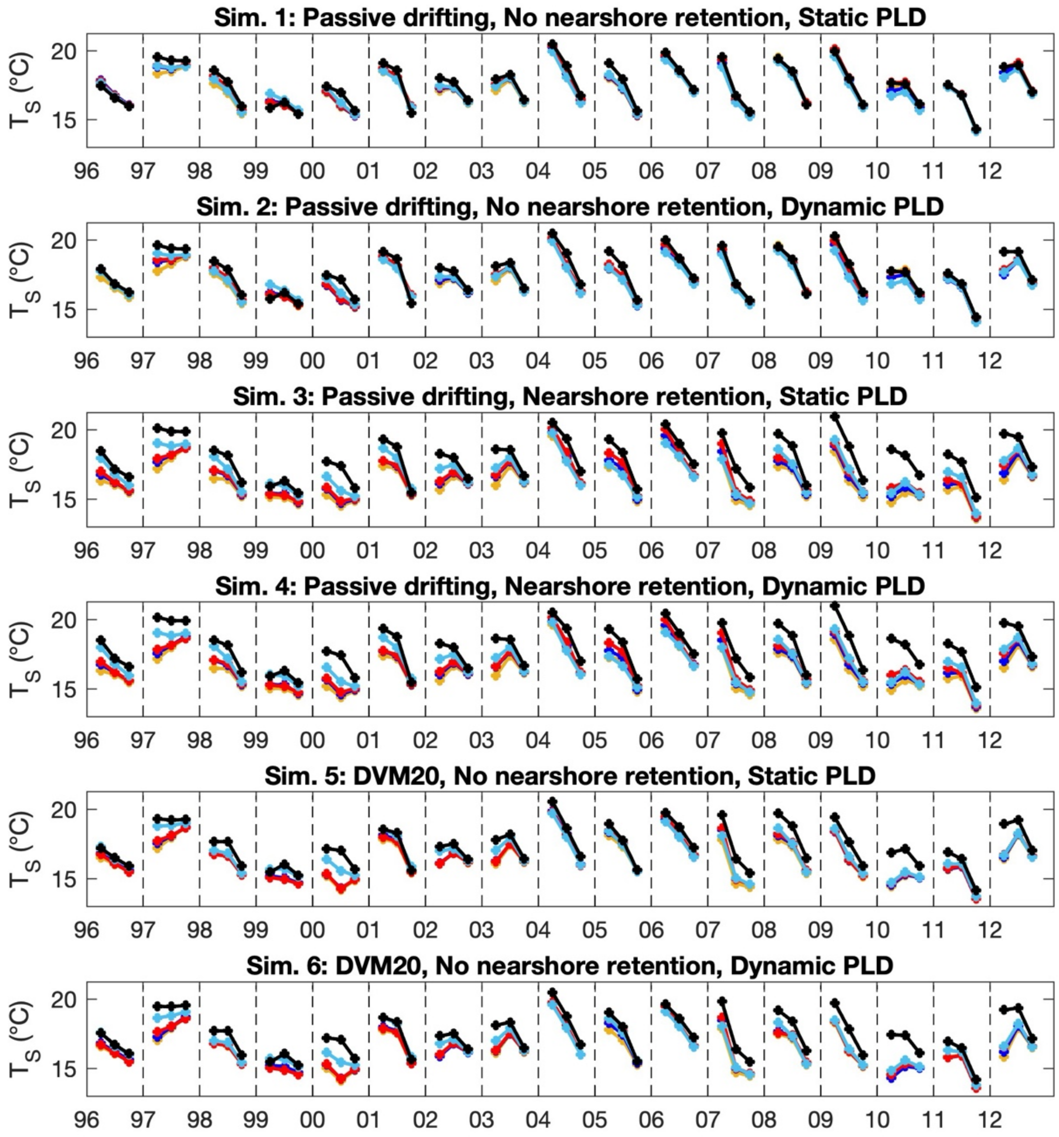


**Figure S11:** Source site temperature ($T_S$,°C) for Simulations 1-6. $T_s$ is shown for the larval reproduction period of September-November. Gaviota is the yellow line. Ellwood is the dark blue line. Stearns is the red line. Anacapa is the light blue line. Scripps is the black line.

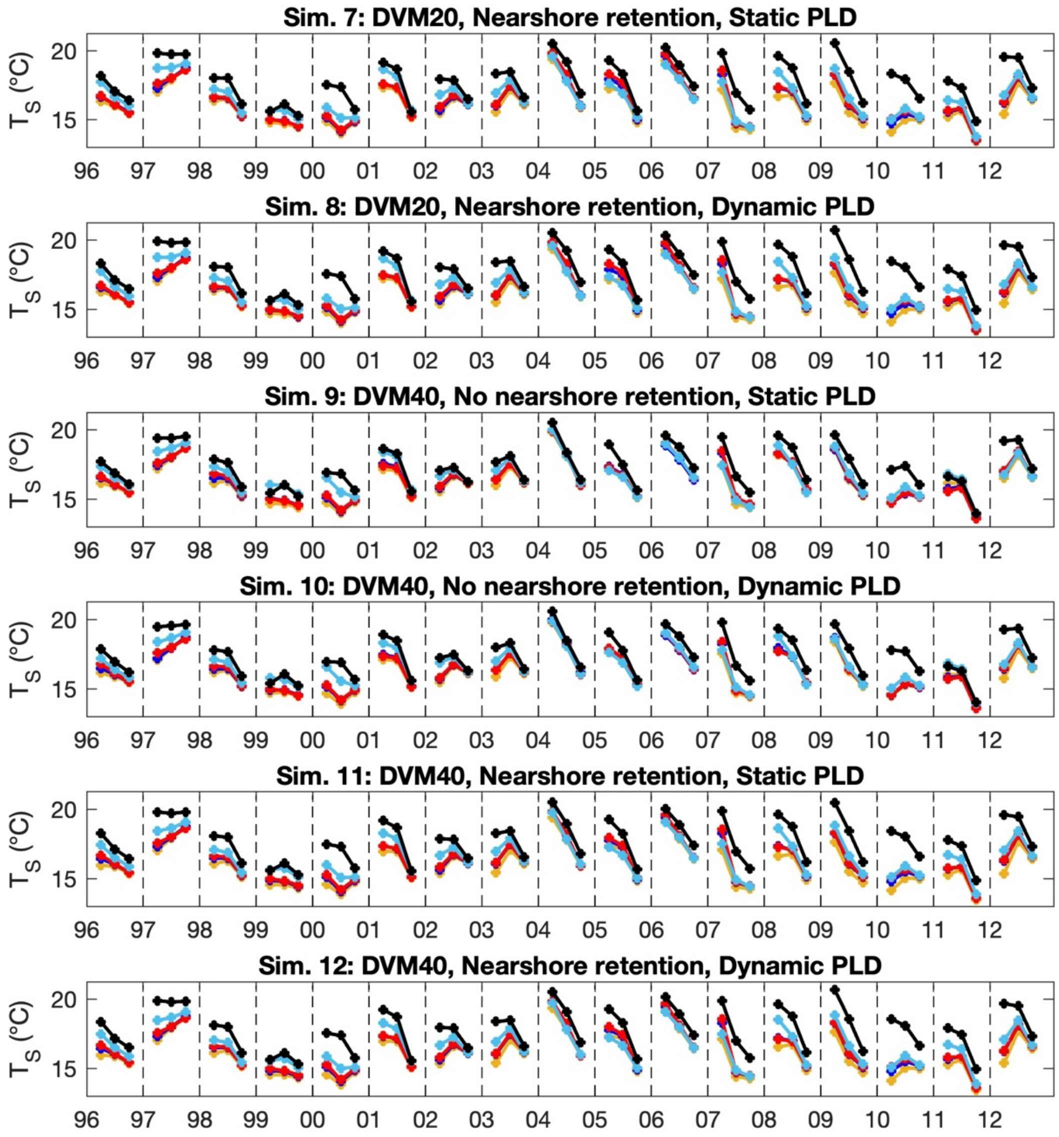


**Figure S11 (con't):** Source site temperature ($T_S$,°C) for Simulations 7-12. $T_s$ is shown for the larval reproduction period of Sept-Nov. Gaviota is the yellow line. Ellwood is the dark blue line. Stearns is the red line. Anacapa is the light blue line. Scripps is the black line.
